# Bioprospecting Novel Luciferase Genes from Museum Coleoptera

**DOI:** 10.64898/2026.04.21.719859

**Authors:** Jack Bate, Patrick Hardinge, Amit P. Jathoul, Michael R. Wilson, James A. H. Murray

## Abstract

Museum collections of Coleoptera contain genetic material of potential interest to biotechnology, and non-destructive DNA extraction enables the preservation of important specimens with concomitant release of mitochondrial and genomic DNA. Mini-barcoding of regions of the mitochondrial cytochrome oxidase subunit I (MT-COI) gene helps identify and eliminate known species from further investigation. Here we identify a novel luciferase gene, using Consensus Degenerate Hybrid Oligonucleotide (CODEHOP) primers targeting the region of the luciferase gene spanning the fourth exon, intron, and fifth exon to detect luciferase gene content and eliminate samples containing known luciferase sequences. Biotinylated luciferase gene probes from the firefly *Photinus pyralis* enabled the enrichment of potential luciferase gene fragments for next-generation sequencing. A bioinformatic analysis suite was then used to identify a luciferase gene sequence from a previously unidentified firefly originally collected in Costa Rica in 2012. We demonstrate that this newly discovered luciferase, termed CRLuc, catalyses a bioluminescent reaction and we determined its emission spectra, Km for the substrates ATP and D-luciferin, and pH stability.

## Introduction

Bioprospecting [30] is the exploration of natural resources for commercially valuable materials such as genetic information or interesting molecules. The world’s museums contain archived specimens of potential interest to bioprospecting including numerous Coleoptera samples. We considered that genetic material from museum fireflies could provide novel luciferase enzymes with possible applications in *in vivo* imaging, ATP sensing for hygiene, and gene assays.

Fireflies belong to the family Lampyridae and are geographically distributed on all continents except Antarctica [8] (BOLD database at www.boldsystems.org). Beetle bioluminescence is also observed in the families Phengodidae and Elateridae, commonly known as glow-worms and click beetles respectively. Evolutionary pressures from bioluminescent courtship displays and aposematism have driven the divergence of the enzyme luciferase responsible for the characteristic bioluminescence. It has been estimated that the luciferase gene ancestor may have diverged approximately 205 million years ago [52] and there are indications that firefly luciferases evolved from a gene duplication event in a non-bioluminescent fatty acyl-coenzyme A synthetase gene [34]. While only about 30 luciferase gene sequences have been derived from firefly species [33], there may be more than 2000 firefly species worldwide [28]. Coleopteran luciferases are of academic and commercial interest due to their high bioluminescence intensity, broad emission colour range and kinetics, all within a standard bioluminescent system.

The best characterised firefly luciferase gene from *Photinus pyralis* has 6 introns and 7 exons, all less than 60 nucleotides long [10] with evidence of a promoter region upstream of the luciferase start codon [9]. The luciferase enzyme from *P. pyralis* is a 62 kilodalton monomer with an N-terminal domain consisting of three sub-domains, and a C-terminal domain as revealed by X-ray crystallography [41] [7]. Luciferase catalysis is a two-stage process of adenylation and oxidation. Firstly, the substrate D-luciferin (LH2) is converted to D-luciferyl adenylate (LH2-AMP) using adenosine triphosphate (ATP). The LH2-AMP is subsequently oxidised in the presence of molecular oxygen to oxyluciferin. Photons are emitted when the excited oxyluciferin returns to the ground energy state [13].

Luciferases are useful tools in various biotechnology applications. Gene expression can be studied by attaching the luciferase gene as a reporter to genes of interest [50]. ATP detection systems for contaminated surfaces utilise luciferase and luciferin to indicate by bioluminescent intensity the extent of bacterial contamination [40] [39] [21]. Bioluminescence imaging (BLI) has become a valuable tool in antiviral research and therapy [27] and new applications are being developed in biomedicine [42]. Novel luciferase enzyme properties and improved characteristics are required to advance new and existing applications, whether these discoveries arise from natural sources or mutagenic exploration by means of rational design and directed evolution. Natural sources of novel luciferase genes include specimens sampled directly from their habitat and, in this study, bioprospecting from museum collections. Natural sources of novel luciferase genes include specimens sampled directly from their habitat and, in this study, bioprospecting from museum collections. Of particular interest would be a shift in spectra for imaging, increased pH tolerance, enzyme thermostability, the bioluminescent intensity, and improved KM values for substrates ATP and D-luciferin.

The recovery of biological material from museum samples requires non-destructive techniques. This has been demonstrated with the extraction of mitochondrial and nuclear DNA from preserved Coleoptera without external morphological damage [12] and has been used successfully on museum samples collected over 200 years ago [43].

This is remarkable because DNA is known to degrade as a function of heat and time [25] and furthermore, specimens were often asphyxiated with chemicals such as ethyl acetate and formalin, causing extensive breaks throughout the genome accelerating degradation independent of age [11]. Unfortunately, the methods of asphyxiation and preservation are rarely recorded for museum samples.

Next generation sequencing techniques such as Illumina require short nucleotide sequences of 100 to 400 base pairs, and the extracted DNA fragment size of historic specimens often falls in this range, making Illumina sequencing appropriate for museum fireflies without the need for sonication. It has been shown that North American firefly genomes are in the range 433 and 2572 million base pairs [26] while the luciferase gene itself is only approximately 2000 base pairs including introns. Recovering sufficient sequence information for this small gene size requires either very deep sequencing of a method or enrichment of the desired sequence relative to all other genome sequences, such as cross-species capture hybridisation [29] of the fragmented genomic DNA extracts. Illumina sequencing reads can then be analysed to provide an assembly of a previously unknown gene through a bioinformatic pipeline.

Here, we describe the PCR amplification of short fragments of the mitochondrial cytochrome c oxidase I (MT-COI) gene using a taxon-specific primer set for the universal amplification of arthropod COI ‘mini-barcodes’, originally designed to amplify a 157 bp region of digestion-degraded DNA extracts from insectivorous bat faecal samples [51]. The target fragment of this primer set lies within the circa 650 bp ‘DNA Barcoding’ region of the COI, and therefore sequencing of the PCR product should enable identification of the species or closest relative where a match was unavailable on the BOLD database. Mini-barcodes lack the equivalent accuracy of the full circa 650 bp region, which provides greater than 97 percent species-level specificity in arthropods [16] but have previously been demonstrated to provide greater than 90 percent species-level resolution in degraded DNA samples [31]. We successfully enriched Illumina libraries from a museum-sourced unidentified firefly, which was sequenced as 150 bp paired-end reads on the Illumina NextSeq 500 Sequencer (Illumina, Inc., CA, USA) in the Cardiff University BIOSI Genomics Research Hub. Bioinformatic analysis was conducted using the Cardiff University School of Biosciences Biocomputing Hub high-throughput computing cluster (YSGO). From the bioinformatics, we were able to identify a novel luciferase from the Costa Rican firefly, which we syntheised and evaluated for potential desirable characteristics. The workflow we describe could be used to target luciferase or other genes from other museum specimens without damage to their external appearance.

To summarise, the unknown Costa Rican firefly was collected in 2012, the North American Firefly Photinus pyralis in 1996 and the British Glow Worm Lampyris noctiluca in 2006. All were DNA extracted to preserve morphology and tested with mini-barcoding to identify the species or closest relative. All three samples were tested with CODEHOP primers to indicate new luciferase sequences and were luciferase gene enriched for sequencing. The fragmented DNA from the unknown Costa Rican firefly was Illumina sequenced and the luciferase gene bioinformatically assembled, the protein synthesised and the enzyme evaluated.

## Materials and Methods

### Materials

All fireflies were obtained from Amgueddfa Cymru – National Museum Wales either from their own collection or from the California Academy of Sciences in San Francisco, via the National Museum Wales, and were loaned with permission to extract genetic material whilst preserving the specimen. The *Lampyris noctiluca* sample was collected in 2006 by Dr Amit Jathoul with permission from private land near Cambridge, UK and cold stored at -80 degrees C in DMSO.

### Non-destructive DNA extraction

Non-destructive DNA extraction buffer [12] (3 mM calcium chloride, 2 percent w/v sodium dodecyl sulphate, 40 mM dithiothreitol, 250 µg/ml proteinase K, 100 mM Tris-HCl pH8, 100 mM sodium chloride) was used to extract DNA from whole Coleoptera. In brief, each specimen was submerged in buffer (300 to 800 µl as required) and incubated for 20 hours at 55 degrees C with gentle agitation (300 rpm). Each beetle was submerged in 100 percent ethanol for 4 hours prior to their return to the Museum collections. Recovered DNA in the buffer was purified by phenol/chloroform/isoamyl alcohol extraction and ethanol precipitation following the standard protocol [38].

### DNA quality and quantification

DNA extracts were quantified on the Qubit fluorometer (Fisher Scientific, Massachusetts, USA) using the dsDNA high sensitivity assay to the manufacturer’s instructions. The fragment size distribution of each DNA extract was calculated using the Tapestation 4200 System (Agilent, California, USA).

### Mini-barcoding

Mini-barcoding is an effective method to distinguish closely related species [18]. Taxon-specific oligonucleotide primers were designed to target a ‘mini-barcode’ region [51] of the mitochondrial cytochrome oxidase subunit I gene (MT-COI) are detailed in Table 1.

**Table 1.**
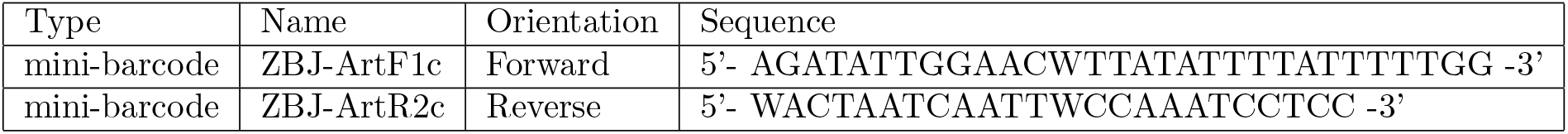
Mini-barcode oligonucleotide primers. Oligonucleotides targeting the mitochondrial cytochrome oxidase subunit I gene for species specific detection.

Primers were synthesised by Sigma-Aldrich (Missouri, USA) and reconstituted to 100 µM with molecular grade water. Primary stocks were diluted to aliquots of 10 µM to reduce freeze-thaw degradation. Mini-barcode PCR used approximately 10 ng of DNA template with 500 nM each primer and 25 µl of Taq PCR Master Mix (Qiagen, Hilden, Germany) (final volume 50 µl) in an Eppendorf Mastercycler (Eppendorf, Hamburg, Germany) following cycling parameters from Zeale et al (2011) [51]. In brief, 95 degrees C for 3 minutes initial denaturation, followed by 16 cycles of: 94 degrees C, 61 degrees C (reducing by 0.5 degrees C per cycle for touchdown), and 72 degrees C; each temperature step being held for 30 seconds, followed by a final 24 cycles with the primer annealing temperature reduced to 52 degrees C and a final 72 degrees C for 10 minutes.

### CODEHOP detection of specific luciferase gene content

Consensus-degenerate hybrid oligonucleotide primers (CODEHOP) [37] [36] [5] are an approach to the amplification of distantly related sequence based on primers designed from consensus protein sequences. Although this approach could not be used to obtain full-length coding sequences due to greater protein sequence divergence at the N and C termini of the protein, we sought to use CODEHOP primers designed against conserved regions of the luciferase gene (Table 2) to confirm whether the samples contained new luciferase sequences.

**Table 2.**
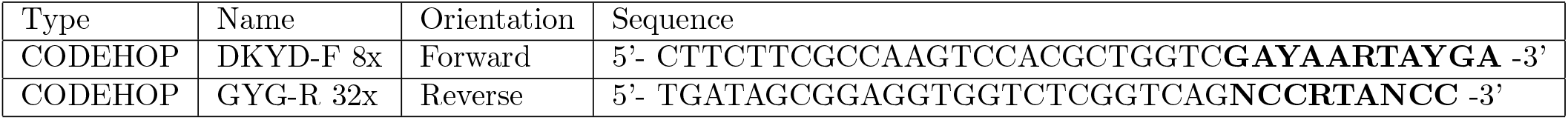
CODEHOP degenerate nucleotide oligonucleotide primers. The forward (DKYD-F) and reverse (GYG-R) CODEHOP primers designed for luciferase gene amplification. 8x and 32x refer to the degeneracy of the degenerate core sequence, shown in bold. Product length is approximately 218 bp.

This set of primers targets the amplification of a region of the luciferase gene that starts in the fourth exon, spans the intervening intron, and ends in the fifth exon. The CODEHOP primer set was initially tested on the DNA extracts of Lnoc and Ppy.

Amplicons were successfully produced, visualised by electrophoresis, and gel extracted by Sanger sequencing. A BLASTn [1] search showed that the sequencing results of both samples matched their respective luciferase genes, with 99.28 percent nucleotide identity match for Lnoc, and 94.67 percent for Ppy. We found that improved resolution of the product bands (SI Figure S2) could be achieved with the supplementation of 4 percent DMSO, without significant inhibition to the amplification in qPCR. From this result, all future use of the CODEHOP primer set was supplemented with 4 percent DMSO. Amplification of the Costa Rica firefly DNA using the same CODEHOP primers was also successful, and the sequencing showed a 86.83 percent nucleotide identity match to the equivalent section of *P. pyralis* luciferase coding region, suggesting a novel luciferase gene sequence (SI Tables S3, S4).

### Luciferase gene enrichment probe

A gBlock synthesised by IDT (Integrated DNA Technologies, Coalville, Iowa, USA) was designed based on mRNA from the *Photinus pyralis* luciferase gene (accession number M15077.1). Terminal primers PpyF and PpyR (Table 3) were used to create a biotin-containing cross-species capture probe using PCR with 50 nM gBlock, 200 nM each primer and 25 µl of Taq PCR Master Mix (Qiagen) with 25 µM Biotin-11-dUTP Solution (Fisher Scientific) (total volume 50 µl) aiming for an approximate inclusion rate of biotin-11-dUTP of 2.5 percent in the final probe.

**Table 3.**
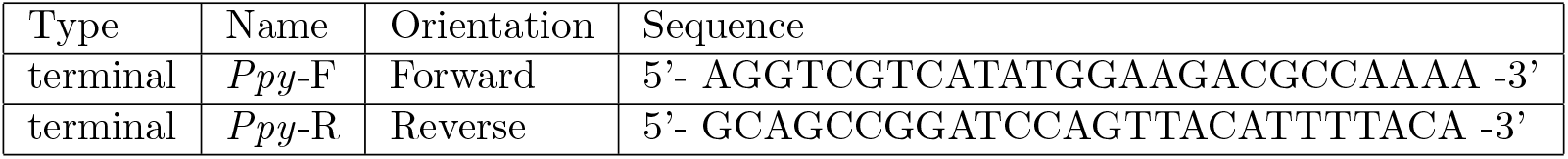
Terminal oligonucleotide primers. The forward (*Ppy*-F) and reverse (*Ppy*-R) terminal primers designed for the luciferase gene enrichment protocol.

In brief, PCR parameters of 94 degrees C for 3 minutes initial denaturation, followed by 30 cycles of 94 degrees C for 40 seconds, 60 degrees C for 40 seconds, and 72 degrees C for 2 minutes followed by a final extension at 72 degrees C for 5 minutes. The biotinylated probe was sized by agarose gel and confirmed to be approximately 1650 base pairs. Multiple reactions were pooled and purified with the QIAquick PCR Purification Kit (Qiagen) following the manufacturer’s instructions.

### Illumina Library enrichment

The NEXTFLEX® Rapid DNA-Seq Kit for Illumina® Platforms (PerkinElmer, Massachusetts, USA) was used to create Illumina sequencing libraries from the firefly genomic DNA extracts. Approximately 1 µg of the Illumina library was mixed with 50 ng of the biotinylated probe and molecular grade water to 7.5 µl. An equal volume of 2X hybridisation solution (1.5 mM sodium chloride, 40 mM sodium phosphate buffer pH7.2, 10 mM EDTA pH8, 10 percent v/v 100X Denhardt’s solution, 0.2 percent w/v sodium dodecyl sulphate) was added and ovelaid with 50 µl of mineral oil (Sigma-Aldrich) and heated to 99 degrees C for 5 minutes. The temperature was lowered from 65 to 55 degrees C at 0.2 degrees C per hour. 100 µl of thoroughly mixed Dynabeads M-280 Streptavidin (Invitrogen, California, USA) were washed three times with 100 µl TEN buffer (10 mM Tris-HCl pH7.5, 1 mM EDTA, 1 M sodium chloride) by magnetic separation and resuspension. 15 µl of the hybridised library was then added to the washed beads in 100 µl of TEN buffer, mixed and agitated at room temperature for 20 minutes. Two low stringency washes (1X SSC buffer 20X, 0.1 percent w/v sodium dodecyl sulphate) at room temperature for 15 minutes each, followed by three high stringency washes (0.1X SSC buffer 20X, 0.1 percent w/v sodium dodecyl sulphate) at 65 degrees C for 15 minutes each, using magnetic separation, resuspension, and agitation. The pelleted beads were eluted with 50 µl freshly prepared 0.1 M sodium hydroxide and agitated for 20 minutes at room temperature. An equal volume of 1 M pH7.5 Tris-HCl was added to the supernatant from magnetic separation and passed through a G50 spin column (Epoch Life Science, Texas, USA) following the manufacturer’s instructions.

### Illumina sequencing

Purified amplified Illumina libraries were sequenced as 150 base pair paired-end reads on the Illumina NextSeq 500 Sequencer (Illumina) in the Cardiff School of Biosciences Genomics Research Hub. Illumina sequencing data was submitted to the NCBI Sequence Read Archive (SRA) and can be accessed using the SRA Run Selector (available at https://www.ncbi.nim.nih.gov/Traces/study/) under the BioProject accession PRJNA802557.

### Bioinformatics for gene assembly

All bioinformatics were processed using Cardiff School of Biosciences Biocomputing Hub computing cluster. The Illumina data was initially processed with Trimmomatic (available at https://github.com/usadellab/Trimmomatic) to perform quality trimming and adapter clipping [4] (see SI Figure S13). Trimmed sequenced data was assessed using FastQC (available at https://www.bioinformatics.babraham.ac.uk/projects/fastqc/) to determine the quality score (see SI Figure S14). The luciferase genes were assembled using a BASH script (SI Figure S15) making use of the packages Bowtie2 [22], SAMtools [24], and SPAdes [2]. Bowtie2 aligns short sequencing reads to larger reference sequences and outputs the alignments in SAM (Sequence Alignment Map). SAMtools provides utilities for post-processing alignments in SAM format, including conversion to BAM (Binary Alignment Map). SPAdes is a de novo assembler and produces contigs (Nodes) from overlapping sequence reads in BAM format. The resulting contigs provide sequence information to determine the luciferase gene of interest.

### Costa Rica firefly luciferase

A gene encoding the proposed Costa Rica Fluc was designed as codon optimised for *E. coli* and was synthesized by IDT and incorporated into a pET16b vector

(Sigma-Aldrich) to enable transformation of *E. coli* BL21 (DE3) competent cells (Fisher Scientific). The colonies arising from transformation were induced with IPTG (Fisher Scientific) and screened with D-luciferin citrate spray as below.

### Bioluminescence detection by Photon Imager Optima

The PhotonIMAGER Optima (Biospace Labs) possesses a photonmultiplier tube (PMT) which enables resolution at a single photon level. The bioluminescent signal is captured and analysed using proprietary software available under license from https://biospacelab.com. 20 second acquisitions were used for luciferin screens and 25 nanometer steps in wavelength for luciferase spectra calculations. All acquisitions were recorded at room temperature. Plated *E. coli* BL21 colonies were transferred using nylon Hybond-N membrane (GE life sciences, Illinois, USA) and placed face up on LB agar plates prepared with 100 µg/ml carbenicillin and previously spread with 200 µl LB broth containing 12.5 µl of 1 M IPTG. The colonies were incubated at room temperature for 3-4 hours. Following induction, the colonies were screened by spraying each plate with 500 µM luciferin in 0.1 M sodium citrate (pH 5) and imaged for bioluminescence.

### Luciferase properties using Clariostar

Bioluminescence emission from purified protein samples were measured in a CLARIOstar Plus Microplate reader (BMG LABTECH, Ortenberg, Germany). The CLARIOstar was preconfigured with the Firefly preset optic setting and the gain set to 2500 across all assays. The following settings were used: Focal height set to 11 millimeters, substrate mix pump volume 100 µl, pump speed 430 µl per second. All acquisitions were performed at room temperature and in triplicate. Data was analysed with MARS software (BMG LABTECH) and further analysed in Microsoft Excel. High resolution biolumnescence spectra between 450 and 800 nanometers (10 nanometer stepwidth) were obtained by injecting the substrate mix onto 50 µl of Fluc to give final concentrations of 1 µM ATP, 500 µM LH2, and 0.167 µM Fluc in 150 µl. Each component was previously diluted in chilled TEM buffer pH7.8. Incubation at room temperature for 30 seconds before acquisition at integrals of 2 seconds. Spectra and flash kinetics were measured at different pH (6.3, 6.8, 7.3, 7.8, 8.3 and 8.8), adjusting with acetic acid or sodium hydroxide. For spectra measurements, incubation at room temperature for 30 seconds preceded acquisition by integrating light emissions over 2 seconds for 36 wavelength scanpoints between 450 and 800 nanometers, with a stepwidth of 10 nanometers. For flash kinetics, no incubation preceded acquisition, with integrals of 20 milliseconds for 1000 consecutive measurements. Peak intensity (Imax) was used to calculate kinetic parameters using the Michaelis-Menten equation [45]. Imax was calculated over a range of substrate concentrations. For ATP this was 0.1, 0.5, 10, 25, 50, 100, 200, 400, 800, and 1000 µM, while maintaining LH2 at 500 µM. For LH2 this was 0.1, 0.5, 1, 5, 10, 20, 35, 70, 140, and 200 µM, while maintaining ATP at 1 mM. On the CLARIOstar injection light emissions were integrated over 20 milliseconds for 1000 consecutive measurements. Imax values were plotted against each substrate concentration. Kinetic constants were calculated from Hanes-Woolf plots [17] [20].

### Analysis tools

Nucleotide sequences were keyword searched and accessed on the European Nucleotide Archive (ENA)(https://www.ebi.ac.uk/ena/home). Basic Local Alignment Search Tool (BLAST) (https://blast.ncbi.nlm.nih.gov) was used to identify nucleotide and protein sequences of high similarity to the novel luciferase gene and protein. Clustal Omega (https://www.ebi.ac.uk/Tools/msa/clustalo) aligns two or more nucleotide or protein sequences. Expasy (https://web.expasy.org/translate) was used to translate nucleotide to amino acid sequences. QIAGEN CLC sequence viewer v8 was used for the construction of nucleotide and protein alignment images.

## Results

### Non-destructive DNA extraction

Non-destructive DNA extraction for Coleoptera [12] was initially attempted on two specimens of Lampyridae (see Materials and Methods) designed to compare DNA quality from a museum specimen and a laboratory specimen stored at -80 degrees C. A dry-preserved museum sample of the North American Firefly *P. pyralis* (Ppy) from the collection of the National Museum of Wales, Cardiff had been collected in 1996 and stored at room temperature, and a specimen of the British Glow Worm *Lampyris noctiluca* (Lnoc) collected by one of the authors (APJ) from a site in Cambridge, UK in 2006 and stored at -80 degrees C in DMSO. Figure 1 provides sample images before and after the non-destructive DNA extraction procedure.

**Figure 1.**
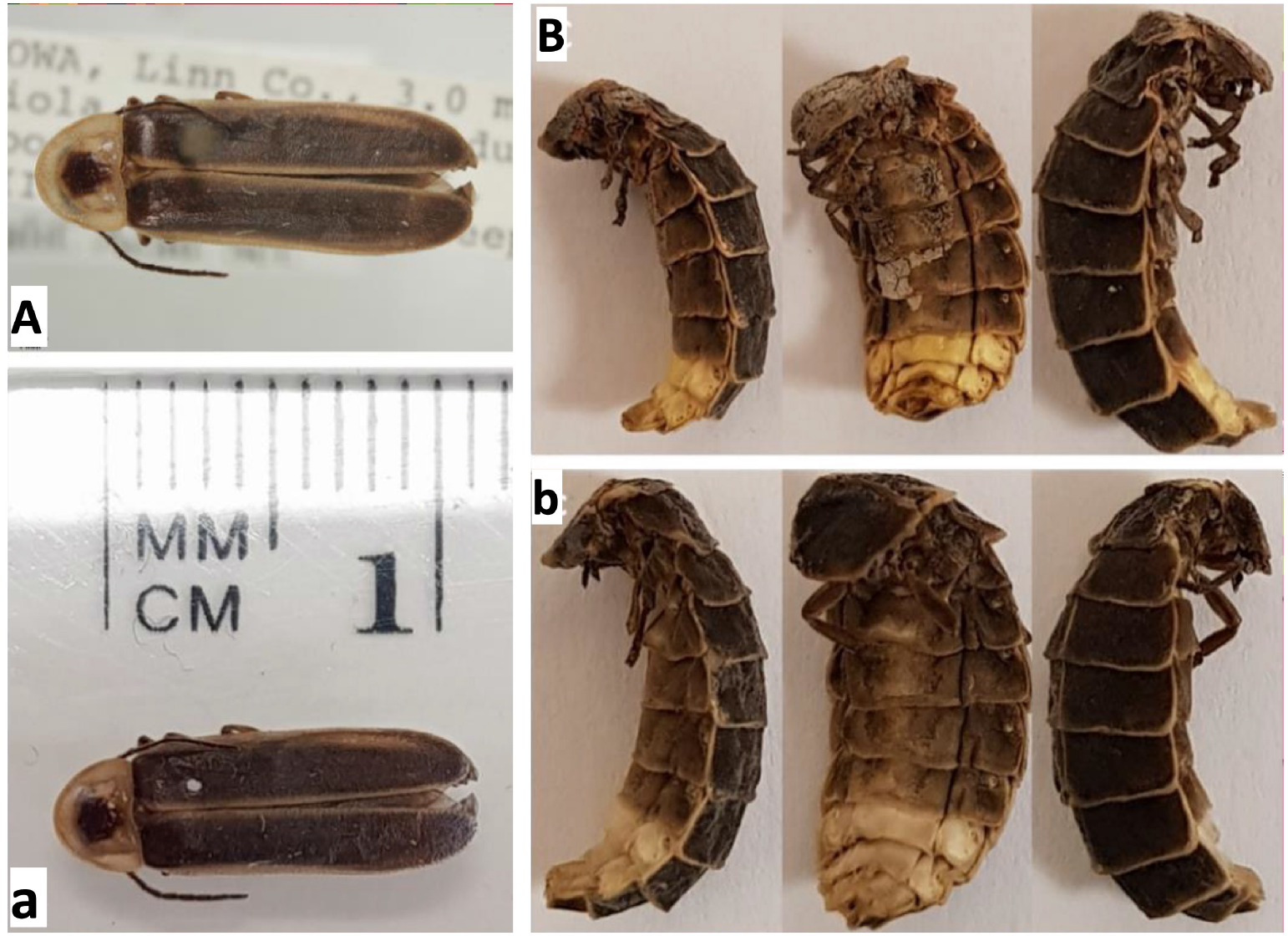
Coleoptera before and after non-destructive DNA extraction. Photographs of *Photinus pyralis* collected in 1996 (A/a) and *Lampyris noctiluca* collected in 2006 (B/b). The images on the top row are the Coleoptera before non-destructive DNA extraction (A/B), and the images on the bottom row are the Coleoptera after non-destructive DNA extraction (a/b). (All photographs courtesy of co-author Jack Bate)

There is limited morphological damage to the specimens; however, some pigment loss from the light organ of the Lnoc sample was noted. This could be attributed to leaching of the pigmented luciferin into the extraction buffer. The museum sample of Ppy is less likely to have luciferin at this level due to sub-optimal storage conditions and would therefore be unaffected by pigment loss. The procedure thus appeared to result in minimal structural changes to the specimens and was then applied to an unknown firefly sample from the National Museum of Wales collected in Costa Rica in 2012, which also displayed only limited morphological damage (SI Figure S1).

### Extraction with DNA fragment size

The quality and quantity of recovered DNA from Ppy and Lnoc samples was assessed by Qubit fluorometry and Tapestation high performance electrophoresis. The dry preserved Ppy recorded a DNA yield of 99.2 ng/µl compared to 164.0 ng/µl recovered from the optimally stored Lnoc sample. Furthermore, the Tapestation results showed the average fragment size to be 17273 base pairs for the Lnoc sample compared to 217 bp for Ppy. This further highlights the role of storage conditions on DNA recovery. We note that the low fragment size from the museum sample would make the feasibility of identifying a complete luciferase gene (approximately 1650 bp of coding sequence, greater than 2000 bp with introns) challenging with PCR. The Costa Rican firefly extraction produced an average DNA fragment size of 881 base pairs but a low DNA concentration of 10.4 ng/µl.

### Mini-barcode method targeting mitochondrial cytochrome c oxidase I gene for species variation

Amplification of a region of the mitochondrial cytochrome oxidase subunit I gene (MT-COI) has previously been demonstrated to provide species-level differentiation with degraded DNA extracts [51]. The Taxon-specific primers designed in that study (Table 1) enable amplification of the extracted DNA for sequencing and searching of the Barcode of Life Database (BOLD), available at www.boldsystems.org [35]. The Lnoc DNA extract was successfully amplified by MT-COI mini-barcode primers. The sequence results from the amplicon provided a 98.11 percent match to the Lnoc mitochondrion partial sequence (accession number MN122858.1). However, amplification with the Ppy extract was not successful for unknown reasons but potentially due to the degree of DNA fragmentation or sequence divergence at the primer binding site. However, Costa Rican firefly DNA was successfully amplified with MT-COI primers and sequencing of the amplicon provided the highest 92.31 percent identity match to the *Photinus australis* MT-COI gene (accession number EU009298.1), suggesting that this may be an individual of a species closely related to *P. australis* [14].

### CODEHOP detection of specific luciferase gene content

Consensus-degenerate hybrid oligonucleotide primers (CODEHOP) [37] [36] [5] are an approach to the amplification of distantly related sequence based on primers designed from consensus protein sequences. Although this approach could not be used to obtain full-length coding sequences due to greater protein sequence divergence at the N and C termini of the protein, we sought to use CODEHOP primers designed against conserved regions of the luciferase gene (Table 2) to confirm whether the samples contained new luciferase sequences. This set of primers targets the amplification of a region of the luciferase gene that starts in the fourth exon, spans the intervening intron, and ends in the fifth exon. The CODEHOP primer set was initially tested on the DNA extracts of Lnoc and Ppy. Amplicons were successfully produced, visualised by electrophoresis, and gel extracted by Sanger sequencing. A BLASTn [1] search showed that the sequencing results of both samples matched their respective luciferase genes, with 99.28 percent nucleotide identity match for Lnoc, and 94.67 percent for Ppy. We found that improved resolution of the product bands (SI Figure S2) could be achieved with the supplementation of 4 percent DMSO, without significant inhibition to the amplification in qPCR. From this result, all future use of the CODEHOP primer set was supplemented with 4 percent DMSO. Amplification of the Costa Rica firefly DNA using the same CODEHOP primers was also successful, and the sequencing showed a 86.83 percent nucleotide identity match to the equivalent section of *P. pyralis* luciferase coding region, suggesting a novel luciferase gene sequence (SI Tables S3, S4).

### Enrichment for sequencing

To obtain the full genomic sequence of the putative luciferases from the total extracted firefly DNA samples, an enrichment strategy for luciferase gene sequences was designed using cross-species capture hybridisation principles. Sequencing libraries were first prepared using the NEXTFLEX® Rapid DNA-Seq Kit (PerkinElmer, Massachusetts, USA) modified to omit sonication in the case of the Ppy due to the already short fragment sizes. Library preparation prior to enrichment was chosen, followed by post-enrichment amplification of the enriched targets, prior to Illumina 2x150 bp sequencing.

Biotinylated probes were synthesised through PCR incorporating Biotin-11-dUTP together with dTTP, achieving an approximate 10 percent biotin-11-dUTP inclusion rate, culminating in probes with about 2.5 percent biotinylated nucleotides as described in Materials and Methods. During the enrichment phase, these probes (Table 3) were combined with the Illumina libraries and subjected to a controlled heating and cooling process allowing for selective hybridisation to sequences of varying specificity (see Methods). Probe-target complexes were then isolated using Dynabeads™ M-280 Streptavidin due to their high affinity to biotin. Magnetic separation then facilitated buffer exchanges to remove unbound non-target sequences, and the targets were eluted using NaOH alkaline denaturation.

Post-capture, the targets were amplified and purified using NEXTFLEX® Rapid DNA-Seq Kit and Agencourt AMPure XP SPRI beads, enhancing target concentration and removing residual probe sequences. Process validation was performed through qPCR with the CODEHOP primer set on both enriched and non-enriched libraries and demonstrated an increase in the relative abundance of luciferase sequence fragments in all enriched libraries relative to the non-enriched equivalents, with the most significant enrichment being observed in the Costa Rica library (Figure 2).

**Figure 2.**
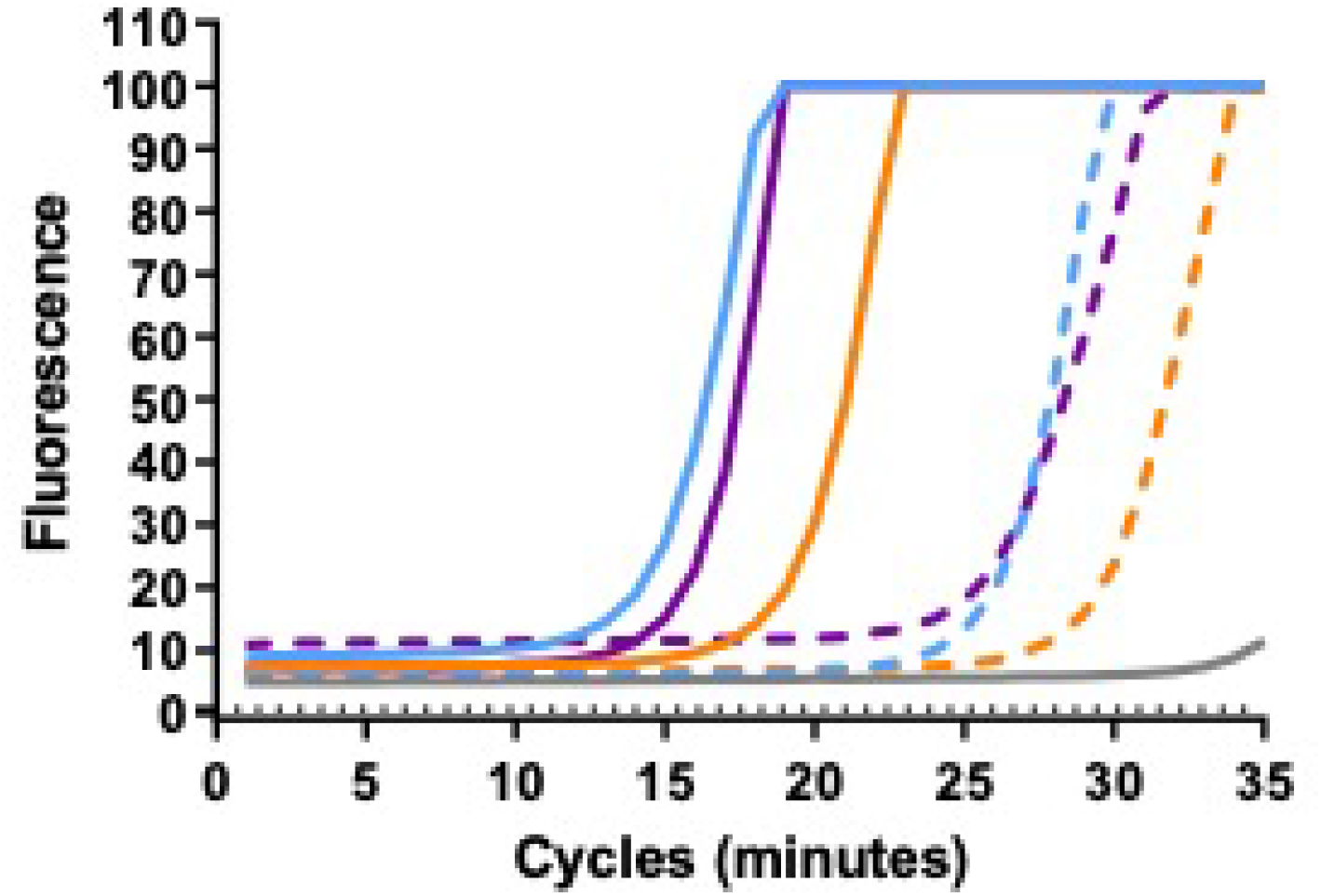
Amplification of libraries before and after enrichment. Quantitative PCR for 35 cycles using CODEHOP primers DKYD-F and GYG-R targetting the luciferase gene. Amplification of enriched libraries of *Lampyris noctiluca* in light blue, *Photinus pyralis* in orange, and the Costa Rican firefly in purple. The amplification of pre-enriched libraries is indicated by dotted lines. Triplicate replicates were used for each and the average values displayed.

Sanger sequencing of the CODEHOP amplification products was used as a final confirmation of novel luciferase gene sequence from the enrichment process, prior to Illumina sequencing. BLAST analysis indicated that all sequences were best matched with *Photinus pyralis*, with the least percent identity being present in the Costa Rica sample at 86.83 percent, whilst the remaining samples ranged between 91.30 – 93.16 percent.

### Sequencing results and bioinformatics

In total, three Nodes were assembled by SPAdes for the enriched Costa Rica, 2012 dataset. The length and coverage of these Nodes are detailed in the Supplementary Information (SI Figures S3 to S9). The three nodes ranged in length from 213 to 1265 bp, and all possessed approximately 90 percent shared identity with voucher specimens of Ppy luciferase complete CDS. Node2 aligned from base 1152 of the Ppy Fluc CDS onward, extending beyond the final base of 2092. The reverse complements of Node1 and Node3 aligned successfully for both Node1RC, which aligned from the start of the reference Ppy Fluc CDS up to base 1228, and Node3RC which aligned between bases 1152 and 1364. Node3RC presents as a region of overlap between the three Node sequences. This common region was aligned and found that although Node1RC and Node3RC were identical, three mismatches with Node2 were present. The first two mismatches occurred in the 4th intron with therefore no influence on the predicted protein sequence. The location of the third mismatch was identified as within the 5th exon and would substitute the 337th codon from CGA to CGC, which both encode arginine, meaning that either of the mismatched bases at this position would not change the final predicted protein sequence. With the construction of a Node consensus complete, an alignment with the cDNA sequence of Ppy Fluc was performed to identify the corresponding regions in the Node consensus which are predicted to comprise the seven exons of the Costa Rica Fluc. These seven predicted exon sequences were extracted and realigned to the Node consensus to visualise the layout of predicted exon and intron sequences in the complete CDS of Costa Rica Fluc. Finally, a translation of the full predicted exon sequence was aligned with the amino acid sequence of Ppy Fluc (Figure **??**), suggesting that the proposed Costa Rica Fluc would be a 550 amino acid protein which shares 93.45 percent amino acid sequence identity with Ppy Fluc (Figure 3).

**Figure 3.**
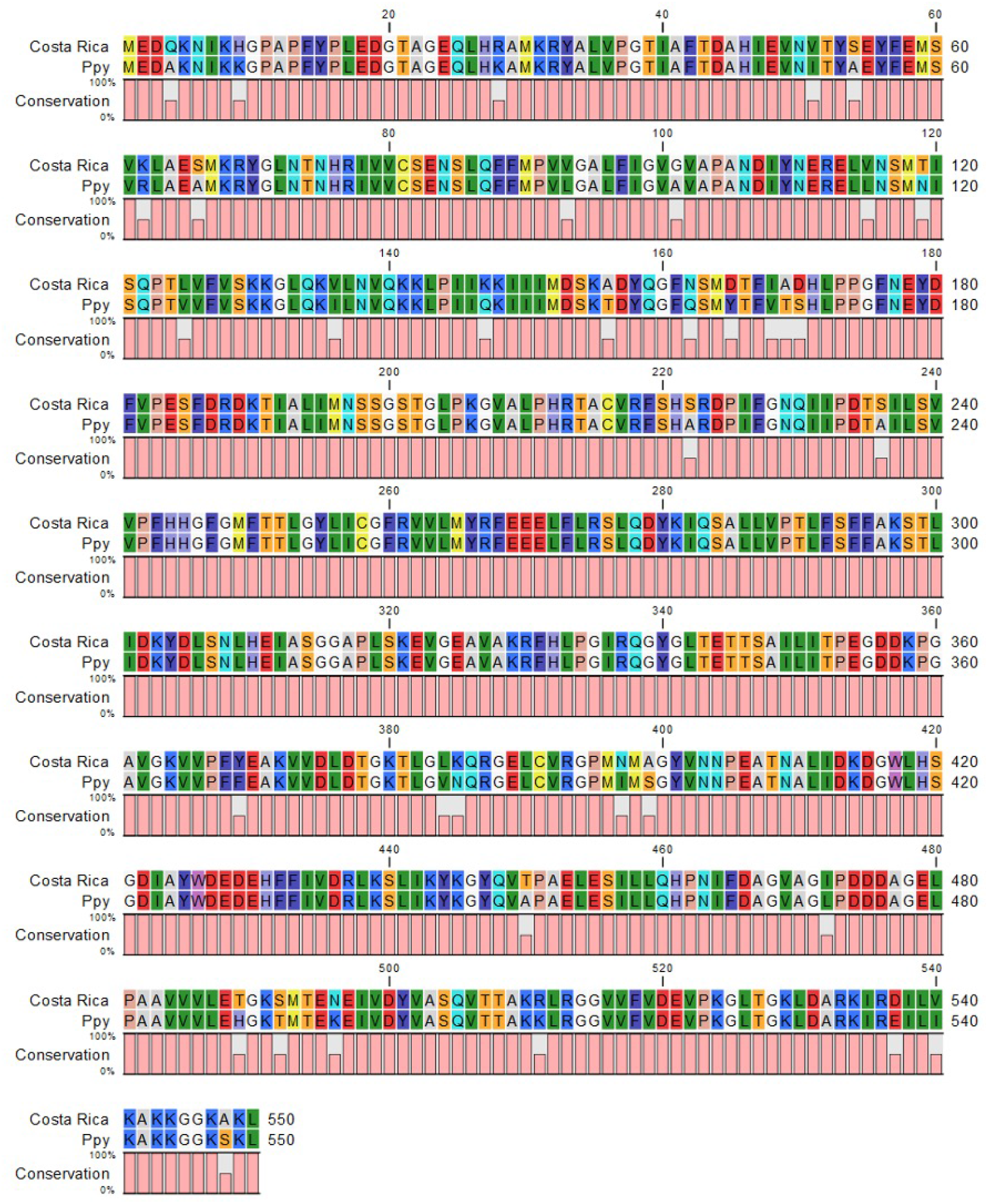
Alignment of Costa Rica Fluc with Ppy Fluc amino acid sequences. Using CLC sequence viewer the proposed translated amino acid sequence of Costa Rica Fluc and the amino acid sequence of Ppy Fluc were aligned with 93.45 percent sequence identity. Mismatches between the two amino acid sequences are indicated by the conservation bar plot in pink.

### Validation and characterisation of novel luciferase CRLuc

With a Costa Rica Fluc protein sequence proposed from the SPAdes output Nodes, functional verification was needed to establish whether the speculative enzyme could catalyse the luciferase reaction and produce a bioluminescence signal. An *E. coli* codon-optimised gene was synthesised and incorporated into the pET16b plasmid to enable transformation of *E. coli* BL21 (DE3) competent cells. The colonies which arose from the transformation were induced for protein production (SI Table S1) with IPTG prior to screening with D-luciferin citrate spray in a PhotonIMAGER Optima.

The bioluminescence data acquired during the screens was analysed in the M3 Vision software package. The bioluminescence signal of the primary screen, as interpreted by M3 Vision, is displayed in Figure 4A.

**Figure 4.**
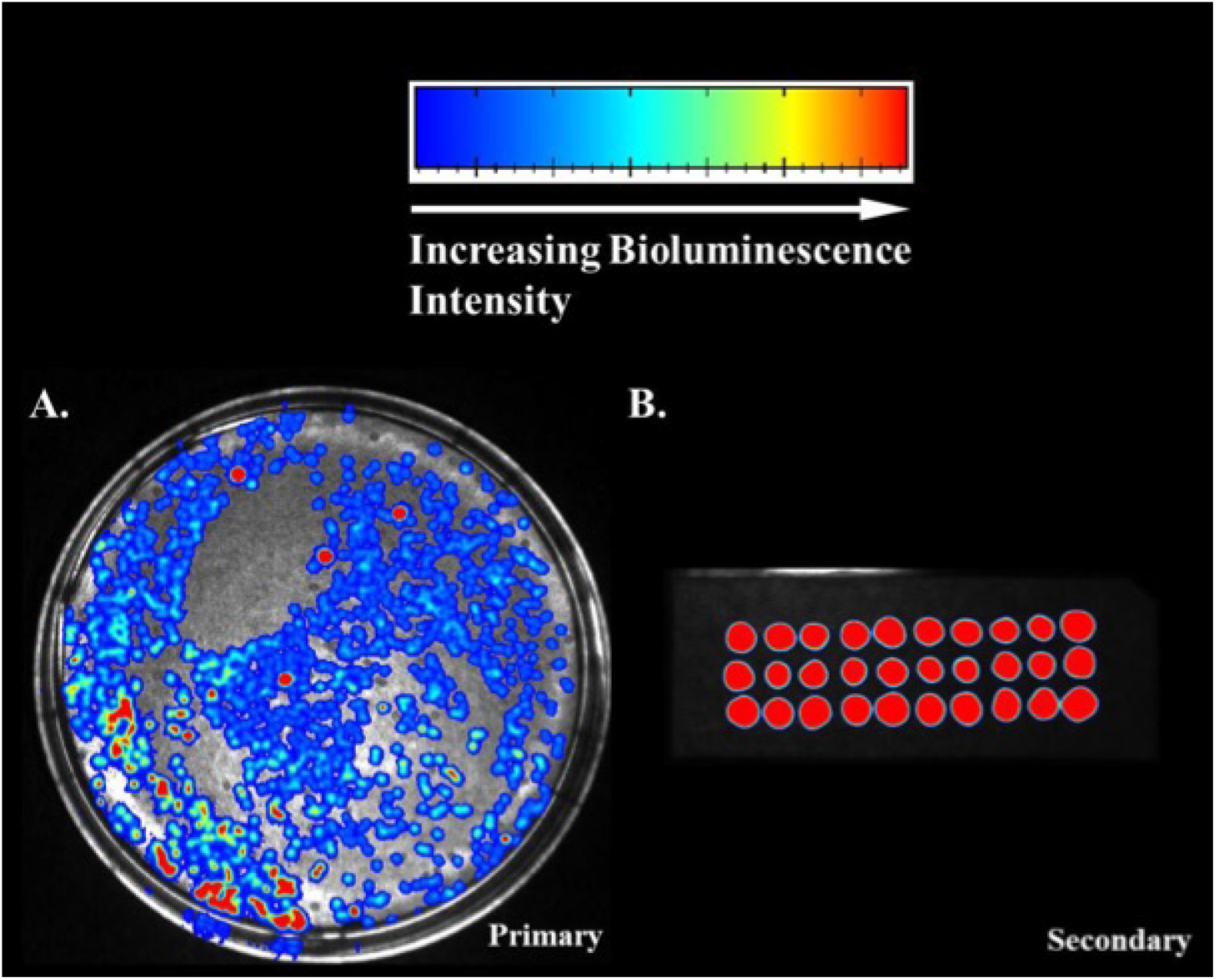
Screening for bioluminescence activity of the Costa Rica Fluc. The plasmid pET16b containing the Costa Rica Fluc gene was used to transform *E. coli* BL21 (DE3) and the colonies were transferred to nitrocellulose, induced with IPTG and screened with luciferin. Bioluminescence was detected on a PhotonIMAGER Optima and the bioluminescent intensity displayed from blue (low intensity) to red (high intensity). The bioluminescent activity of the plated transformants from the primary screen over a 20 second intergral is shown in (A). In (B) randomly picked colonies from the primary screen, were screened for bioluminescent activity. Images were produced using M3 Vision software and the intensity scaling between (A) and (B) is not directly comparable.

A secondary screen was conducted using colonies picked from this plate and is shown in Figure 4B. Both the primary and secondary screen confirmed that the proposed Costa Rica Fluc was a functioning coleopteran luciferase, capable of producing a strong bioluminescence signal.

High resolution bioluminescence spectra of purified enzymes were measured using the CLARIOstar Plus Microplate reader which is equipped with a monochromator that allows for the collection of spectra with a resolution of up to 1 nanometer. The bioluminescence spectra lambda max for the control enzyme Ppy Fluc was circa 558 nanometers (Figure 5, SI Figures S10, S11) which is in broad agreement with the recorded lambda max for this enzyme from available literature [23] and thus were taken to indicated that the spectral properties recorded for the remaining enzymes were similarly accurate.

**Figure 5.**
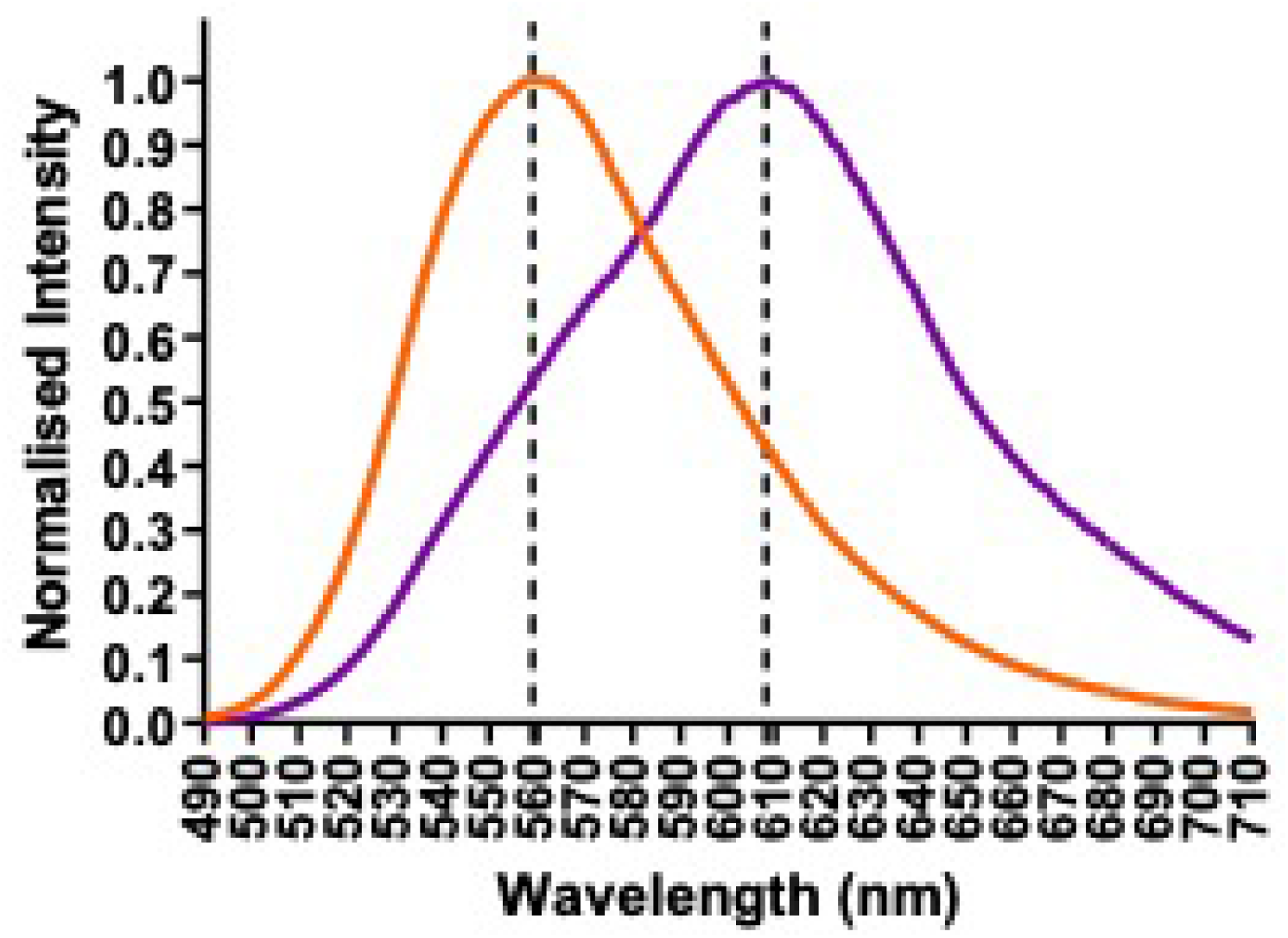
Bioluminescence spectra of Ppy and CRLuc with Luciferin. Lumi-nometric measurements by injection of substrate mix onto each enzyme solution prior to acquisition of bioluminescence. Light emissions between 490 and 710 nanometers, with increments of 1 nanometer for integrals of 2 seconds. PPy Fluc spectra displayed in orange, Costa Rica Fluc in purple. The mean of triplicate results was normalised by relative intensity to lambda max.

The spectra of CRLuc over the pH range 6.3 to 8.8 exhibited a red-shifted emission peak of circa 609 nanometers with a broad bandwidth of 95 nanometers FWHM extending toward the green region (Figure 6).

**Figure 6.**
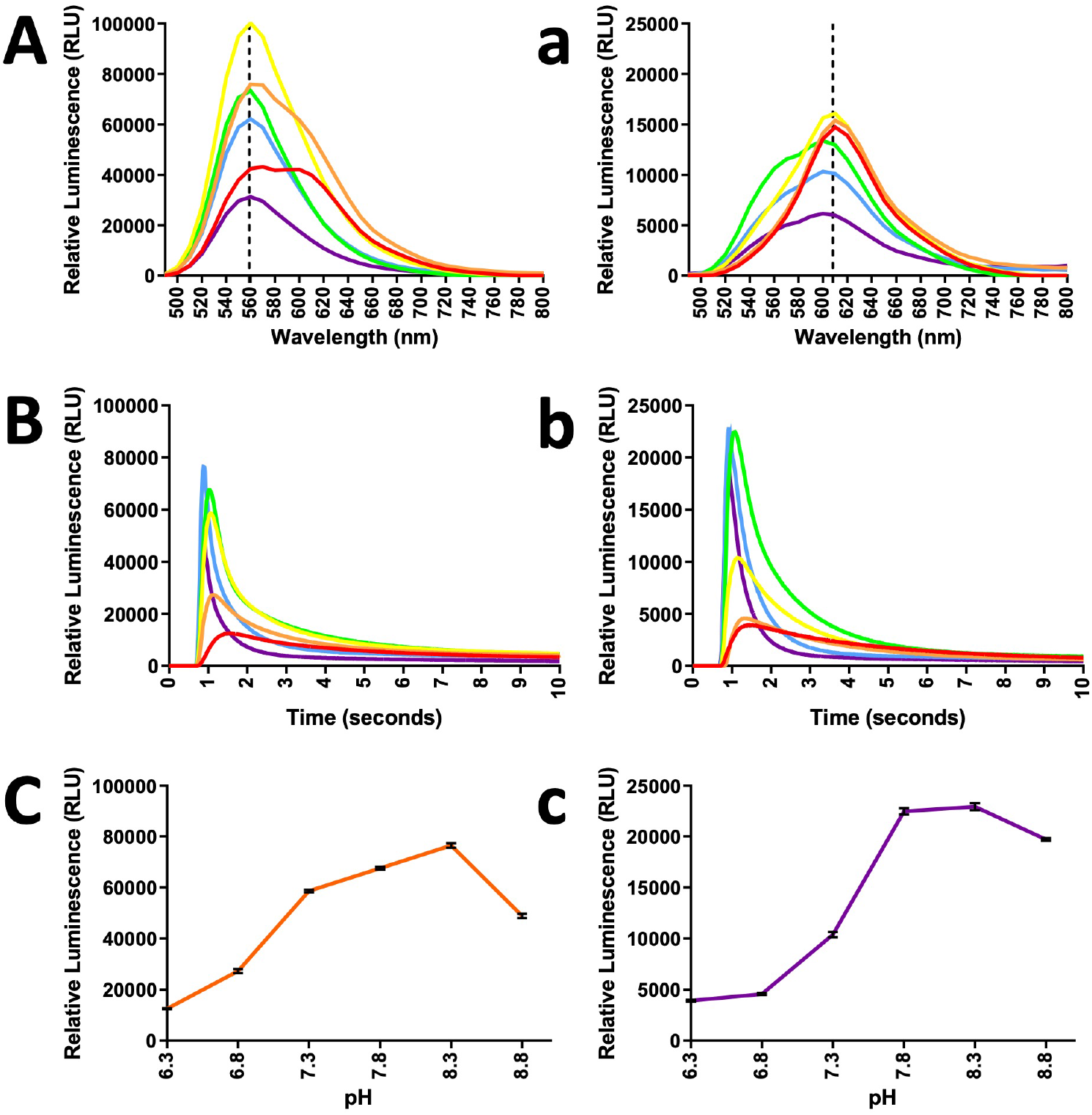
pH dependence of bioluminescence spectra, activity and intensity. Luminometric measurements by injection of substrate mix onto each enzyme such that final assay concentrations were equal to 1 millimolar ATP, 500 micromolar luciferin, and 0.167 micromolar protein. (Aa) bioluminescent spectra, (Bb) bioluminescent activity at pH 6.3 (red), 6.8 (orange), 7.3 (yellow), 7.8 (green), 8.3 (blue) and 8.8 (purple). Light intensity for Ppy Fluc (C) in orange and CRLuc (c) in purple.

The expression of luciferase with red shifted spectra in combination with a standard Ppy Fluc or blue shifted luciferase would enable studies of multiple gene expression. The bioluminescence intensity over the range of pH conditions (6.3 to 8.8) were similar for Ppy Fluc and CrLuc (Figure 6) with a slight shift to higher pH for the CrLuc. However, the light intensity values are lower for CrLuc across the pH range. Tolerance to higher and lower pH and across a wide range of pH would be beneficial to applications such as ATP detection on contaminated surfaces.

With regards to Ppy Fluc, the KM range for LH2 reported in previous studies extends 10 to 20 µM [6] [47] [44], although it has also been recorded as low as 6.6 µM for a recombinant enzyme from Promega [23]. Here, the purified sample of Ppy Fluc displayed a KM for LH2 of 5 µM, and for ATP this value was determined as 45 µM (SI Table S2), in contrast to the range of 56 to 250 µM that has previously been reported [19] [46] [6] [49]. The bioprospected Costa Rican firefly luciferase, CRLuc, was shown to have a higher KM for both substrates than the values here reported for Ppy Fluc, with 14.29 µM calculated for LH2 and 81.11 µM for ATP. These CRLuc KM values do however align with the Ppy Fluc ranges reported elsewhere [23], suggesting that the substrate affinities of both enzymes are comparable (Figure 7 and SI Figure S12). It is desirable to have low KM values for luciferin and ATP which would indicate that the luciferase has a high affinity for its substrates.

**Figure 7.**
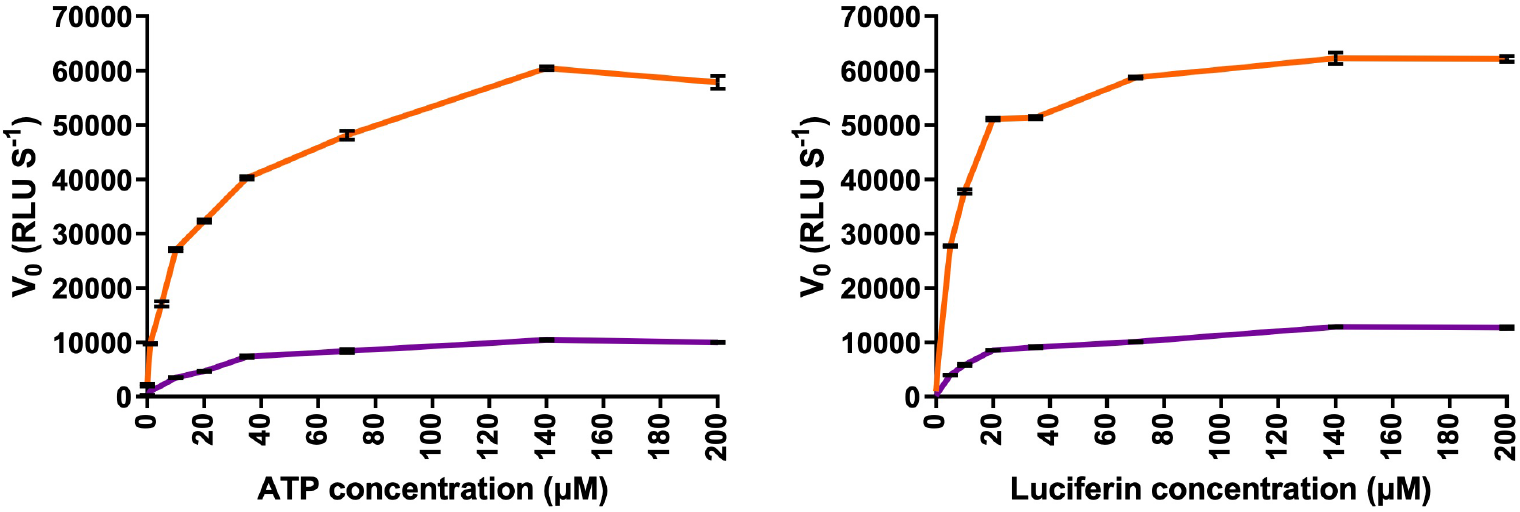
Michaelis-Menten plots of PPy Fluc and Costa Rica Fluc for substrates D-luciferin and ATP. Luminometric measurements of substrate mix injected into each enzyme mix. The substrate D-luciferin is displayed in the left panel with PPy Fluc in orange and Costa Rica Fluc in purple. The right panel shows the results from the substrate ATP. All assays were performed in triplicate and average flash-height measurements (I_max_) are used as an approximation of initial velocity (v_0_).

## Discussion

The pharmaceutical industry has been the largest beneficiary of bioprospecting but other industries such as agriculture, manufacturing and cosmetics have also benefitted [3]. The practice is not without controversy and a legal framework called the Nagoya Protocol on Access and Benefit Sharing (ABS) [32] was put in place to ensure fair and equitable sharing of these benefits. Under the agreement, the utilization of genetic resources should contribute to the sustainable use of biodiversity [15] (https://www.gov.uk/guidance/abs). Although the protocol came into force in October 2014, compliance measures require ratification by the country of origin and country of benefit. Biological resources before this date are not covered by the convention.

The non-destructive DNA extraction process [12] was effective in preserving the appearance of the *Photinus pyralis* and *Lampyris noctiluca* samples (Figure **??**), however the fragility of the Costa Rican firefly was evident (SI Figure S1). The quality of the extracted DNA varied and there appeared to be correlation between specimen age and storage conditions. The dry-preserved Ppy collected in 1996 had an average fragment size of 217 bp, whereas the Lnoc sample collected in 2006 and stored fresh at -80 degrees C had an average size of 17273 bp. Attempts to extract from earlier collected specimens were less successful and may be due to the use of chemical asphyxiates such as ethyl acetate, which was commonly used without consideration for future genetic analysis [11]. Without knowing which chemical asphyxiates were used it was not possible to use age of specimen as an indicator of the quality of DNA available for extraction as the double stranded break mechanism from chemical asphyxiates will accelerate degradation regardless of the age of the museum specimen. The average fragment size of DNA extracted from the Costa Rican firefly specimen collected in 2012 was 881 bp. This may indicate that the specimen was not dispatched with a chemical asphyxiate and the degradation observed was from age and the temperature and humidity of the storage conditions. Further refinement of the methods described here could however allow greater use of older samples with chemically damaged DNA.

The origin and region of greatest Lampyrid diversity is presumed to be Tropical America; however, the diversity remains largely unknown. The BOLD database (www.boldsystems.org) contains 177 species with barcodes in the family Lampyridae. The largest genus of Lampyridae in the Americas is *Photinus* with over 235 identified [48] and the BOLD database holds 486 *Photinus* specimen barcodes from 28 species. We used mini-barcoding to PCR amplify small fragment targets to enable BLAST to identify highest identity matches as a percentage. This method has previously been demonstrated on degraded DNA samples to provide ¿90 percent species-level resolution [31]. A match between a known luciferase gene sequence and the sequence from an unidentified museum firefly would be discounted for further investigation. The mini-barcode investigation of the Costa Rican firefly showed a 92.31 percent sequence identity with the mitochondrial COI region of the North American firefly *Photinus australis*, a species closely related to *P. pyralis* from individuals collected from Indiana to Florida, USA, and first described by Green (1956) [14]. This indicates that the Costa Rican firefly specimen is likely to be a previously unidentified firefly closely related to *Photinus australis* and was therefore worth further study. For the effective determination of phylogenetic relationships from barcoding to understand Lampyridae diversity, a considerable expansion in the number of species and samples would be required. Therefore the Costa Rican firefly specimen is assumed to be previously unidentified.

We developed a CODEHOP approach to detect the presence of novel luciferase gene sequences in the DNA extract libraries of unidentified museum fireflies. The technique was also used to highlight whether nuclear DNA targets could be amplified as well as the demonstrated mini-barcoding amplification of the mitochondrial COI DNA target. Initially we demonstrated successfully CODEHOP amplification in trial libraries of Ppy and Lnoc. In museum firefly libraries we identified a conserved region of 49 bp in our amplification sequences which could not be matched directly to the Ppy luciferase gene and therefore probe contamination could be excluded. The amplification product from the pre-enriched Costa Rican firefly library was successfully sequenced and matched the sequence from the enriched Costa Rican firefly library CODEHOP amplification results, excluding some terminal variation and a single mismatch. The 49 bp conserved region and the full Costa Rican firefly CODEHOP amplification sequences were aligned to the completed CDS assembled enriched Costa Rican firefly library. Shared identity between the CDS and the full CODEHOP amplification was 79.65 percent and 89.90 percent for the conserved region. This suggests that these amplifications are not of the Costa Rican firefly luciferase gene sequence. The use of 4 percent DMSO with the CODEHOP primers did, however, amplify the Costa Rican firefly luciferase gene with an 86.93 percent identity match to Ppy luciferase.

The bioinformatic analysis of the trimmed sequence reads showed that the quality was good using FastQC and mapping was successful for the enriched Costa Rican firefly library. Using *Photinus pyralis* as a reference sequence, a gene sequence was determined with sequence identity of 88.79 percent to Ppy luciferase complete CDS. It was noted that no reads were able to map to the reference genomes of the Japanese firefly *Aquatica lateralis* (formerly *Luciola*) and the bioluminescent click beetle and Carribean native *Ignelater luminosus* and the sequence identity to the luciferase gene sequence for *Aquatica lateralis* (GenBank: Z49891.1) was only 60.38 percent (there is no luciferase gene sequence available for *Ignelater luminosus*) which could be explained by divergence in sequence identity of these luciferase genes. Those reads that successfully mapped to the Ppy Fluc gene (SI Figure S7) had greater than 88 percent sequence identity and therefore were sufficiently complementary to exceed the unknown threshold. Bowtie2 mapping was shown to be required before assembly with SPAdes for the Costa Rican firefly library. We also demonstrated that the enrichment strategies was essential to the success of the Costa Rican firefly library. The principle of cross-species affinity enrichment has previously been demonstrated to enrich sequences 10 to 13 percent divergent from the probe identity [29] and our Costa Rican firefly results support this. The use of multiple firefly luciferase genes to produce a larger pool of probe sequences would potentially increase the success of enrichment.

The assembler SPAdes produced three Nodes (SI Figures S3, S4, S5) which overlapped to create a single 2304 bp contig. Three mismatches were observed in the 213 bp overlapping region (SI Figure S6) relating to the 4th intron and a synonymous substitution at the 337th codon, all of which are inconsequential to the Costa Rica Fluc. However, these mismatches in a relatively short section could indicate that further errors could be present in the derived Costa Rica gene sequence which may not be inconsequential. The luciferase gene sequence can therefore only be described as representative of the true gene. Using the exon-intron structure of Ppy Fluc for comparison, we were able to determine the exon sequences which shared a 93.45 percent amino acid sequence identity with Ppy Fluc. Nevertheless we demonstrate that our predicted sequence encodes a novel and functional luciferase enzyme with comparable activity to Ppy Fluc.

## Conclusion

We have demonstrated that dry-preserved Museum Coleoptera collections are a potentially valuable source of genetic material. The use of mini-barcoding is simple and targeted; however, it is limited by the availability of barcode sequences of Lampyridae species to enable identification of divergent genes. The CODEHOP method, although useful, could be improved by increasing the pool of divergent gene probes. However, cross-species affinity enrichment with an appropriate bioinformatic pipeline shows the greatest potential for degraded DNA samples. Illumina sequencing is appropriate for the short DNA fragments from non-destructive DNA extraction. Using this method resulted in a novel luciferase gene sequence from an unidentified Costa Rican firefly which encoded a luciferase enzyme capable of bioluminescent functionality. Whilst this enzyme cannot be definitively claimed as an accurate representation of the wild-type enzyme in nature, it serves as a novel source of luciferase gene variation, which is yet to be meaningfully characterised. The unidentified Costa Rican firefly was collected in 2012; much older fireflies are present in museum collections, and this, therefore, is a limitation of the study.

## Supporting Information

A separate supplementary information file is available. The details of the supplementary figures and tables are listed below.

**S1 Figure**

**Costa Rican Firefly post non-destructive DNA extraction**. Photographs of unknown Costa Rican firefly after DNA extraction (photograph taken by Jack Bate).

**S2 Figure**

**Optimisation of CODEHOP primers with DMSO. (A)** Amplification of Lnoc gDNA by quantitative PCR with CODEHOP primers DKYD-F and GYG-R supplemented with DMSO concentrations between 0 and 10 percent. Average of triplicate results plotted with results truncated to greater than 20 cycles. NTCs for each DMSO concentration. **(B)** Amplicons from **(A)** visualised by electrophoresis with NTCs to the left of each positive sample.

**S3 Figure**

**Node2 alignment with Ppy coding sequence**. The alignment of Node2 from the SPAdes assembly of the Costa Rica luciferase gene with the complete CDS of the Ppy Fluc gene. Mismatches between the two sequences are highlighted in grey.

**S4 Figure**

**Reverse complemented Node1 alignment with Ppy coding sequence**. The alignment of Node1RC from the SPAdes assembly of the Costa Rica luciferase gene with the complete CDS of the Ppy Fluc gene. Mismatches between the two sequences are highlighted in grey.

**S5 Figure**

**Reverse complemented Node3 alignment with Ppy coding sequence**. The alignment of Node3RC from the SPAdes assembly of the Costa Rica luciferase gene with the complete CDS of the Ppy Fluc gene. Mismatches between the two sequences are highlighted in grey.

**S6 Figure**

**Three mismatches in the alignment of Node1RC, Node2 and Node3RC**. The alignment of all three Nodes from the SPAdes assembly of the Costa Rica luciferase gene cropped to the length of Node3 sequence. The three mismatches between the two sequences are highlighted in grey.

**S7 Figure**

**Node consensus alignment with Ppy coding sequence**. The alignment of the consensus Node contigs of the Costa Rica luciferase gene with the complete CDS of the Ppy Fluc gene. Mismatches between the two sequences are highlighted in grey. Sequence identity 88.79 percent.

**S8 Figure**

**Node consensus alignment with Ppy luciferase gene cDNA**. The alignment of the consensus Node contigs of the Costa Rica luciferase gene with the cDNA sequence of the Ppy Fluc gene. The seven exon regions of the two sequences are highlighted in blue.

**S9 Figure**

**Node consensus alignment with predicted exons for CRLuc**. The alignment of the consensus Node contigs of the Costa Rica luciferase gene with the predicted exon sequences of the Costa Rica luciferase. The seven exon regions of the two sequences are highlighted in red.

**S10 Figure**

**Normalised bioluminescence spectra against pH for Ppy Fluc and CRLuc with LH2**. Luminometric measurements by substrate injection to each enzyme after 30 seconds RT incubation. Light emission from 2 second integrals every 10 nanometers from 450 to 800 nanometers. Average of triplicate values shown with data normalised with light intensity relative to lambda max.

**S11 Figure**

**Normalised bioluminescence against pH for Ppy Fluc and CRLuc with LH2 for 20 seconds**. Luminometric measurements by substrate injection to each enzyme. Light emission from 20 millisecond integrals for 1000 measurements. Average of triplicate values shown for each pH with data normalised with light intensity relative to emission peak.

**S12 Figure**

**Hanes-Woolf plots of Ppy Fluc and CRLuc with LH2 and ATP**. Plots of [S]/v against [S], where [S] is the substrate concentration and v is the estimated initial rate at that concentration indicated by Imax. Triplicate measurements of Imax over substrate concentration ranges for ATP and LH2 were used to calculate average kinetic parameters. In all assays the final concentration of enzyme was estimated to be 0.167 micromolar. For kinetic parameter calculations for LH2, assays were performed with final concentrations between 0.1 and 200 micromolar, saturated with 1 millimolar ATP. Kinetic parameters of ATP measurements, assays were performed with final concentrations between 0.1 and 1000 micromolar, saturated with 500 micromolar LH2.

**S13 Figure**

**BASH script for Trimmomatic with annotations**. Script to process sequenced libraries with Trimmomatic (Bolger et al. 2014). Annotations that describe the function of the following section of code are shown in blue. Sample names and their designation of ‘$*{*i}’ are shown in purple to indicate where in the script each sample name will be substituted. The script is looped to process each sample. R1 and R2 files are forward and reverse paired reads, respectively. Singletons are reads which have no identified pairing. Trimmomatic is available at https://github.com/usadellab/Trimmomatic.

**S14 Figure**

**BASH script for FastQC with annotations**. Script to generate quality reports on trimmed libraries with FastQC. Annotations that describe the function of the following section of code are shown in blue. Sample names and their designation of ‘$*{*i}’ are shown in purple to indicate where in the script each sample name will be substituted. The script is looped to process each sample. R1 and R2 files are forward and reverse paired reads, respectively. Singletons are reads which have no identified pairing. FastQC is available at https://www.bioinformatics.babraham.ac.uk/projects/fastqc/.

**S15 Figure**

**BASH script for luciferase gene extraction with annotations**. Script to extract reads corresponding to luciferase gene sequences from trimmed libraries through the execution of Bowtie2 (Langmead and Salzberg 2012), SAMtools (Li et al. 2009), and SPAdes (Bankevich et al. 2012). Annotations that describe the function of the following section of code are shown in blue. Sample names and their designation of ‘$*{*i}’ are shown in purple to indicate where in the script each sample name will be substituted. The script is looped to process each sample. Similarly, reference genomes and their designation of ‘$*{*j}’ are shown in red to indicate where in the script each reference genome name will be substituted. Again, the script is looped to process each sample. R1 and R2 files are forward and reverse paired reads, respectively. Singletons are reads which have no identified pairing. Bowtie2 is available at https://sourceforge.net/projects/bowtie-bio/. SAMtools is available at http://samtools.sourceforge.net. SPAdes is available at https://github.com/ablab/spades. Firefly reference genomes Alat1.4, Ppyr1.4, and Ilumi1.3 are available at http://fireflybase.org/.

**S1 Table**

**Average protein concentrations of two luciferases**. The concentration of Ppy Fluc and CRLuc as determined by Bradford assay of desalted purified protein. Values for SDS-PAGE determined by ImageJ software. The protein sizes are calculated from the protein sequences. Micromolar concentrations are highlighted in grey.

**S2 Table**

**Summary of kinetic parameters for Ppy Fluc and CRLuc**. Average kinetic parameter as derived from triplicate measurements of Imax across a substrate concentration range.

**S3 Table**

**Luciferase genes used in CODEHOP design**. Eleven coleopteran luciferase genes used in the design of CODEHOP primers DKYD-F and GYG-R.

**S4 Table**

**Sequencing from CODEHOP DKYD-F ¿ GYG-R amplification**. Details of sequencing results for amplifications with CODEHOP primers DKYD-F and GYG-R. The description, accession, and percent identity of the top BLAST match are provided.

**S5 Table**

**NGS data accession details**. Individual accessions of the Costa Rica firefly dataset under the BioProject accession PRJNA802557. Accessions are made available for access using the NCBI SRA Run Selector (available at https://www.ncbi.nlm.nih.gov/Traces/study/), NCBI BioSample (available at https://www.ncbi.nlm.nih.gov/biosample), and an overview of experiment details at NCBI SRA (available at https://www.ncbi.nlm.nih.gov/sra).

## Data availability

Information on the data underpinning the results presented here, including how to access them, can be found in the Cardiff University data catalogue at http://doi.org/10.17035/cardiff.27044365. Illumina sequencing data was submitted to the NCBI Sequence Read Archive (SRA) and can be accessed using the SRA Run Selector (available at https://www.ncbi.nlm.nih.gov/Traces/study/) under the BioProject accession PRJNA802557.

## Acknowledgments

The authors would like to express their gratitude to the Cardiff University Genomics Hub for their assistance with sequencing.

## Author contributions

JAHM, PH, APJ and JB conceived the experiment(s), MRW supplied materials and guidance, JB conducted the experiment(s), JB analysed the results and discussed the findings with JAHM, PH, APJ and MRW. All authors reviewed the manuscript.

