## Supplementary Information for "Bioprospecting Novel Luciferase Genes from Museum Coleoptera"

**Authors list**

Jack Bate^1,2^, Patrick Hardinge^1*^, Amit P Jathoul^1,3^, Michael R Wilson^4^, and James A H Murray^1^

**Affiliation**

^1^School of Biosciences, Cardiff University, Museum Avenue, Cardiff CF10 3AX, UK
^2^Cultech Ltd., Unit 2 Christchurch Road, Baglan Industrial Estate, Port Talbot SA12 7BZ, UK
^3^Bioflares Ltd., Building 500, Discovery Park, Ramsgate Road, Sandwich CT13 9FF, UK
^4^Department of Natural Sciences, National Museum of Wales, Cardiff CF10 3NP, UK

*

**Supplementary materials**

**Figures**

Supplementary Figure S1. Costa Rican Firefly post non-destructive DNA extraction.

Supplementary Figure S2. Optimisation of CODEHOP primers with DMSO.

Supplementary Figure S3. Node2 alignment with Ppy coding sequence.

Supplementary Figure S4. Reverse complemented Node1 alignment with Ppy coding sequence.

Supplementary Figure S5. Reverse complemented Node3 alignment with Ppy coding sequence.

Supplementary Figure S6. Three mismatches in the alignment of Node1RC, Node2 and Node3RC.

Supplementary Figure S7. Node consensus alignment with Ppy coding sequence.

Supplementary Figure S8. Node consensus alignment with Ppy luciferase gene cDNA.

Supplementary Figure S9. Node consensus alignment with predicted exons for CRLuc.

Supplementary Figure S10. Normalised bioluminescence spectra against pH for Ppy Fluc and CRLuc with LH2.

Supplementary Figure S11. Normalised bioluminescence against pH for Ppy Fluc and CRLuc with LH2 for 20 seconds.

Supplementary Figure S12. Hanes-Woolf plots of Ppy Fluc and CRLuc with LH2 and ATP.

Supplementary Figure S13. BASH script for Trimmomatic with annotations

Supplementary Figure S14. BASH script for FastQC with annotations

Supplementary Figure S15. BASH script for luciferase gene extraction with annotations.

**Tables**

Supplementary Table S1. Average protein concentrations of two luciferases.

Supplementary Table S2. Summary of kinetic parameters for Ppy Fluc and CRLuc.

Supplementary Table S3. Luciferase genes used in CODEHOP design.

Supplementary Table S4. Sequencing from CODEHOP DKYD-F > GYG-R amplification.

Supplementary Table S5. NGS data accession details.

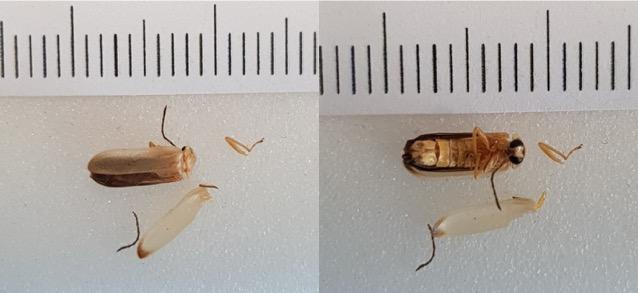

**SI Figure S1: Costa Rican Firefly post non-destructive DNA extraction.** Photographs of unknown Costa Rican firefly after DNA extraction (photograph taken by Jack Bate).

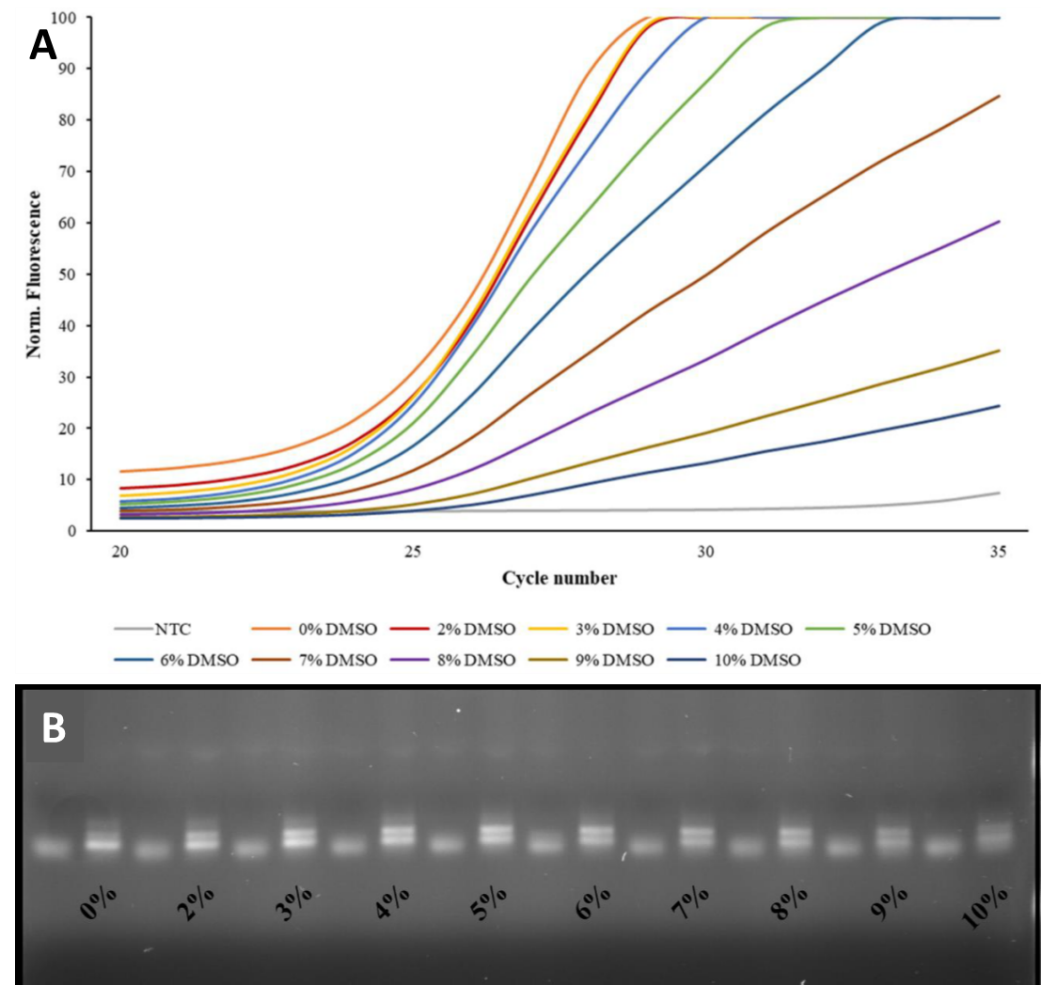

**SI Figure S2: Optimisation of CODEHOP primers with DMSO.** (**A**) Amplification of Lnoc gDNA by quantitative PCR with CODEHOP primers DKYD-F and GYG-R supplemented with DMSO concentrations between 0 and 10 percent. Average of triplicate results plotted with results truncated to greater than 20 cycles. NTCs for each DMSO concentration. (**B**) Amplicons from (**A**) visualised by electrophoresis with NTCs to the left of each positive sample.

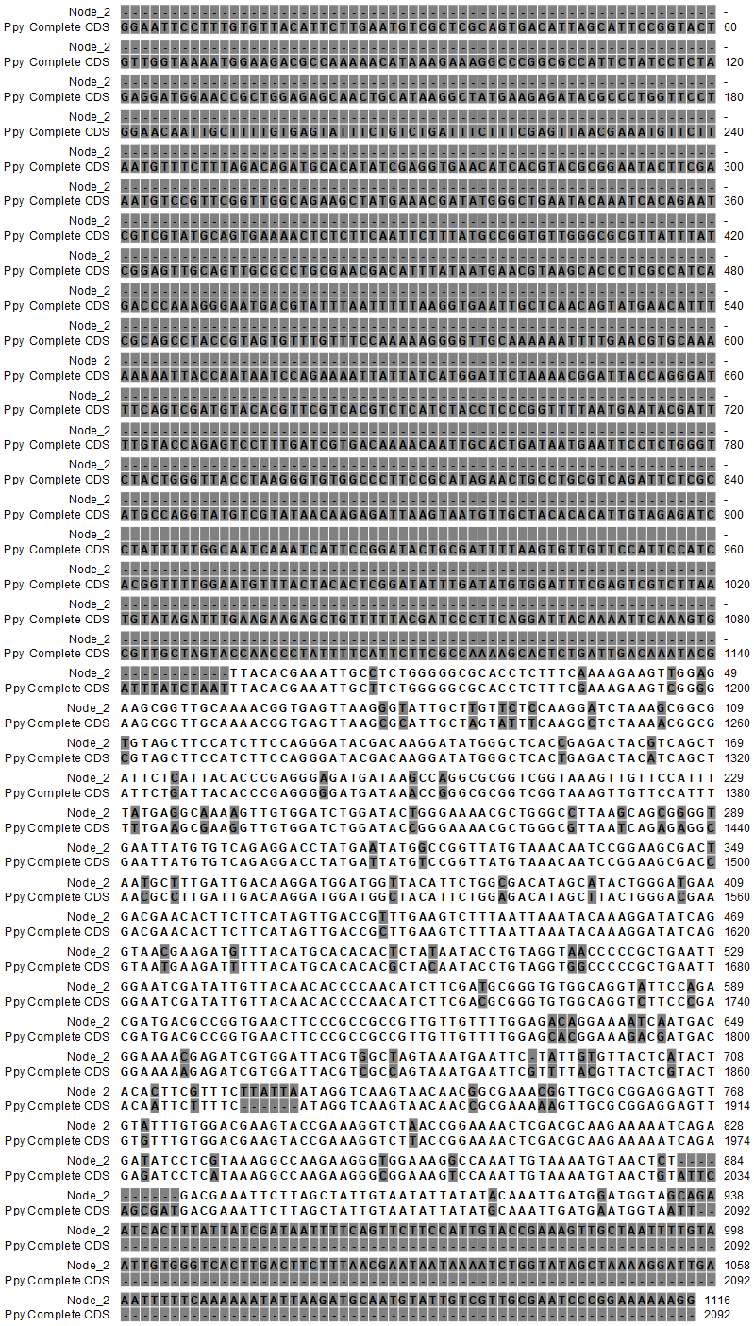

**SI Figure S3: Node2 alignment with Ppy coding sequence.** The alignment of Node2 from the SPAdes assembly of the Costa Rica luciferase gene with the complete CDS of the Ppy Fluc gene. Mismatches between the two sequences are highlighted in grey.

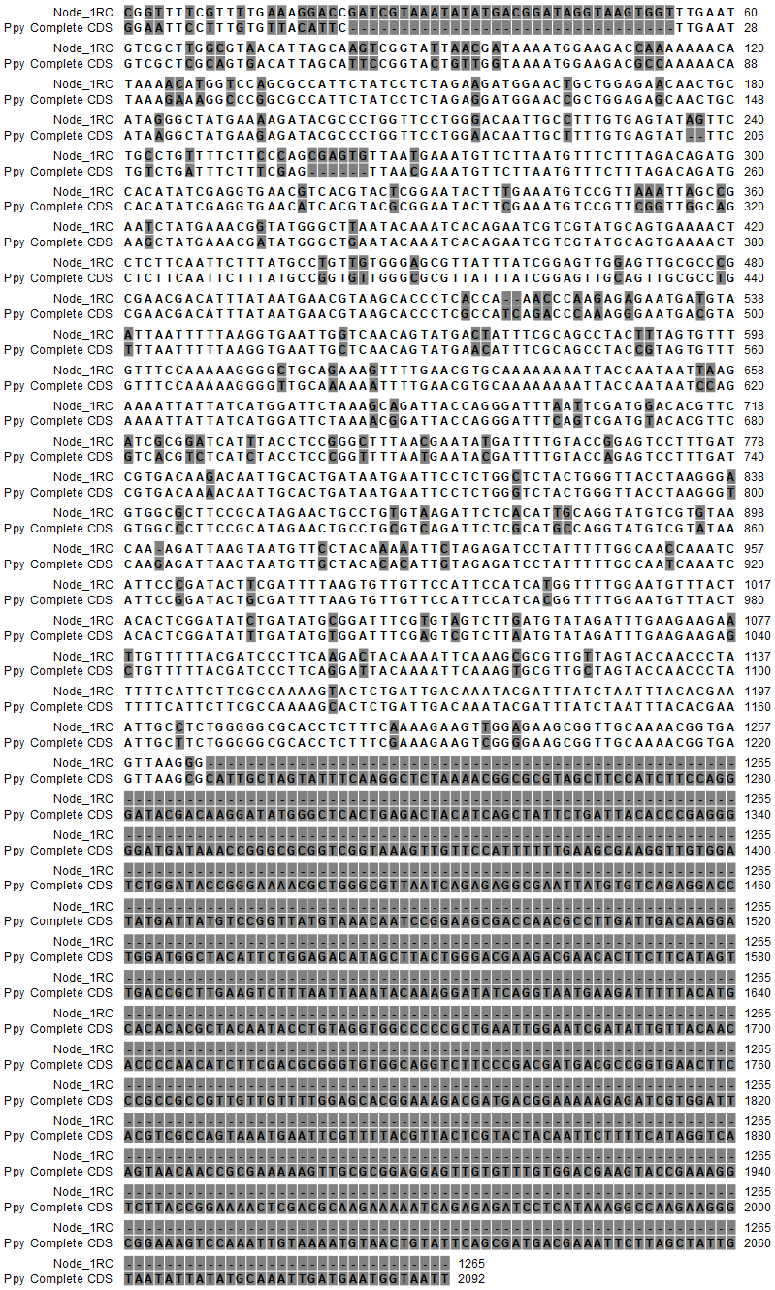

**SI Figure S4: Reverse complemented Node1 alignment with Ppy coding sequence.** The alignment of Node1RC from the SPAdes assembly of the Costa Rica luciferase gene with the complete CDS of the Ppy Fluc gene. Mismatches between the two sequences are highlighted in grey.

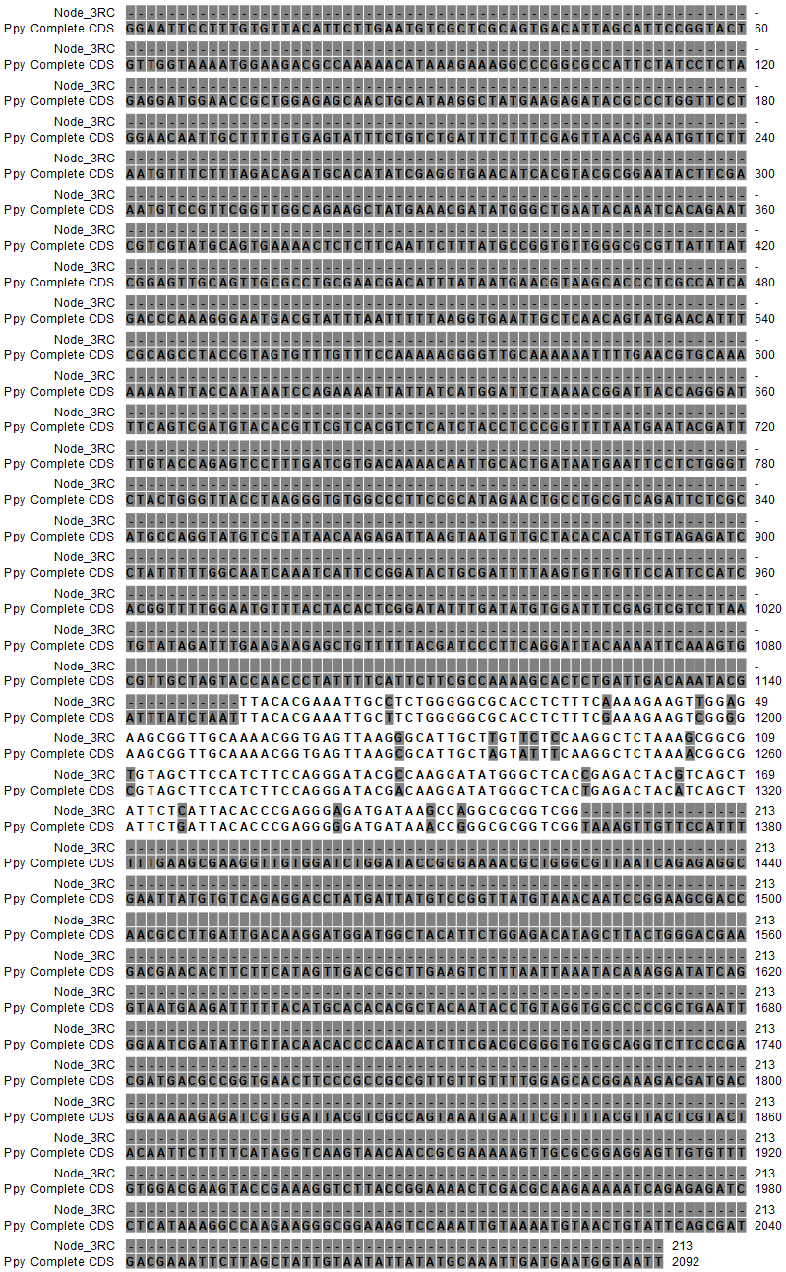

**SI Figure S5: Reverse complemented Node3 alignment with Ppy coding sequence.** The alignment of Node3RC from the SPAdes assembly of the Costa Rica luciferase gene with the complete CDS of the Ppy Fluc gene. Mismatches between the two sequences are highlighted in grey.

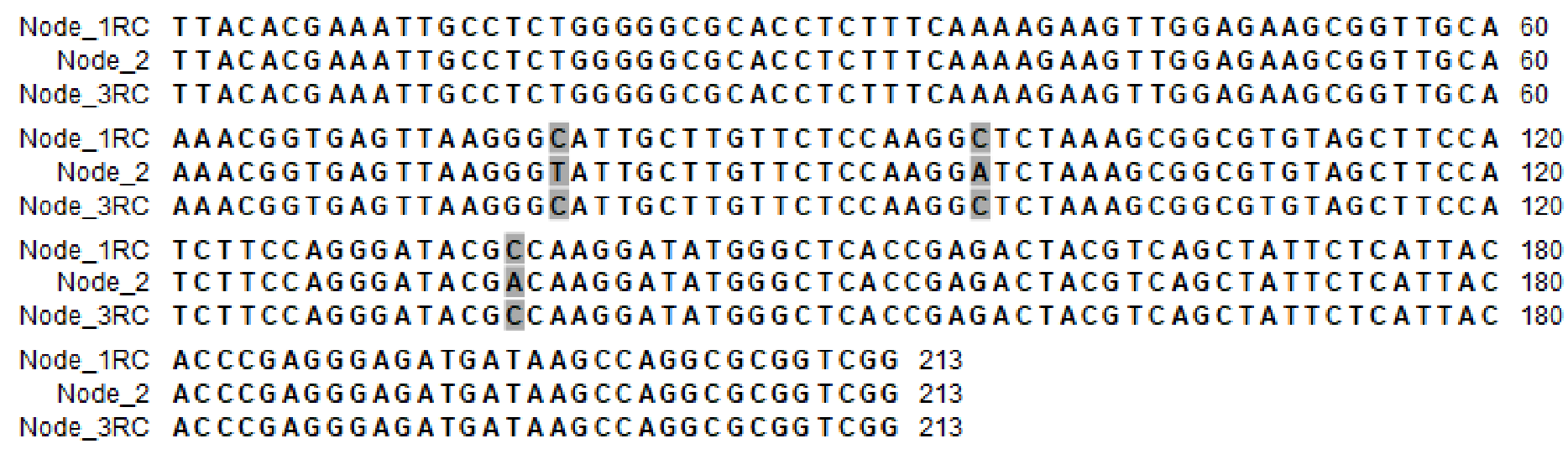

**SI Figure S6: Three mismatches in the alignment of Node1RC, Node2 and Node3RC.** The alignment of all three Nodes from the SPAdes assembly of the Costa Rica luciferase gene cropped to the length of Node3 sequence. The three mismatches between the two sequences are highlighted in grey.

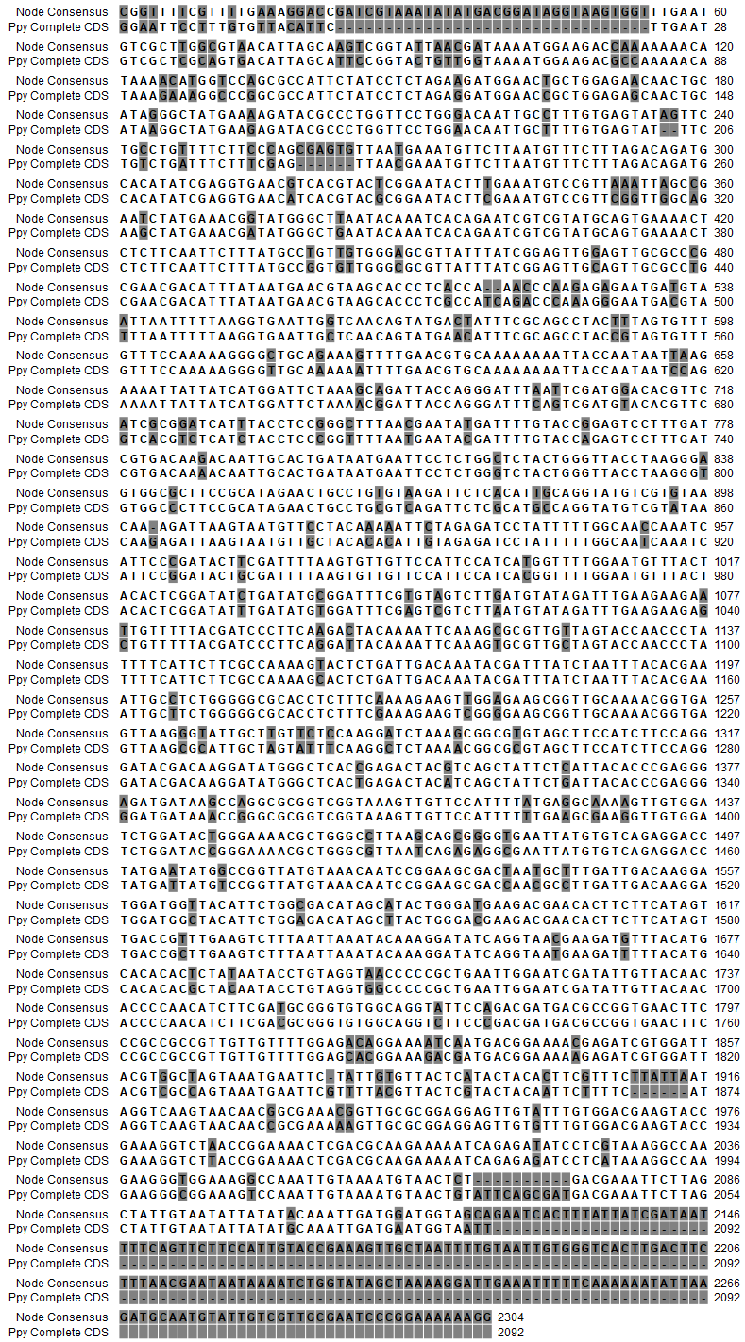

**SI Figure S7: Node consensus alignment with Ppy coding sequence.** The alignment of the consensus Node contigs of the Costa Rica luciferase gene with the complete CDS of the Ppy Fluc gene. Mismatches between the two sequences are highlighted in grey. Sequence identity 88.79 percent.

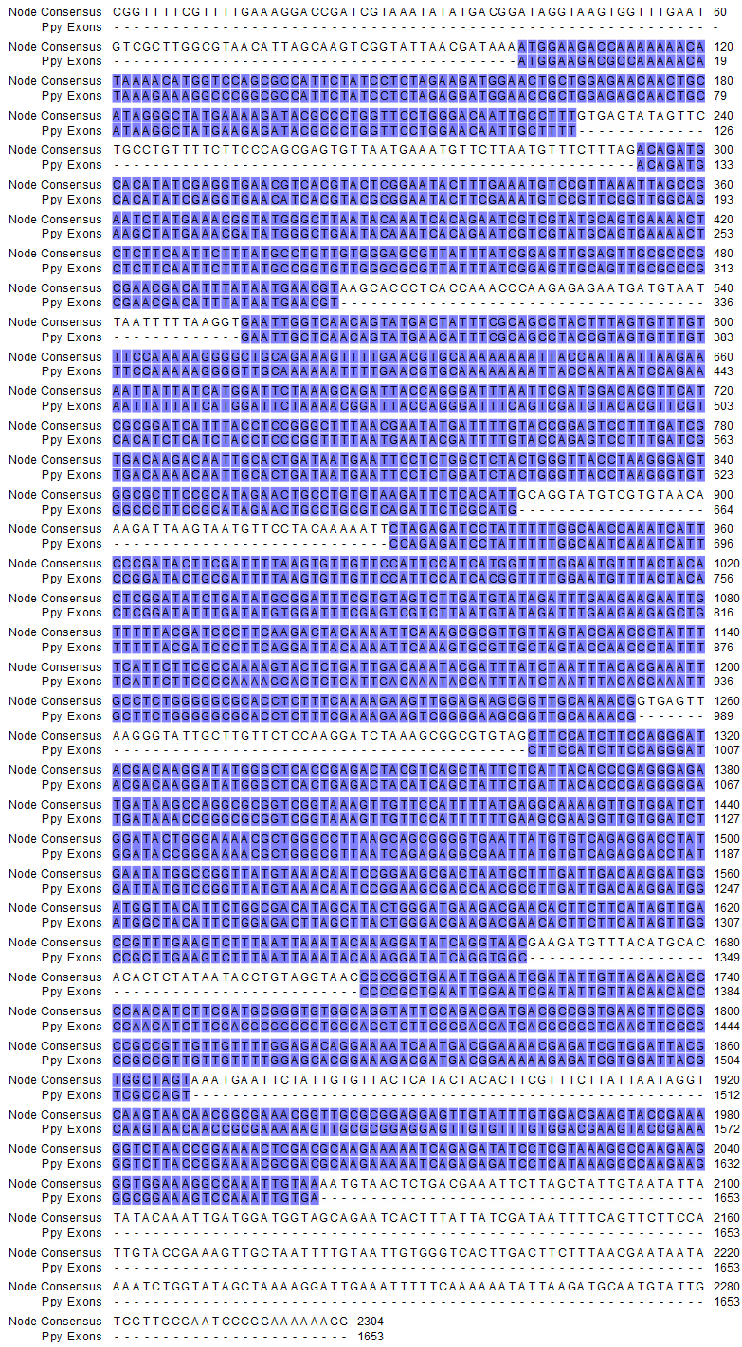

**SI Figure S8: Node consensus alignment with Ppy luciferase gene cDNA.** The alignment of the consensus Node contigs of the Costa Rica luciferase gene with the cDNA sequence of the Ppy Fluc gene. The seven exon regions of the two sequences are highlighted in blue.

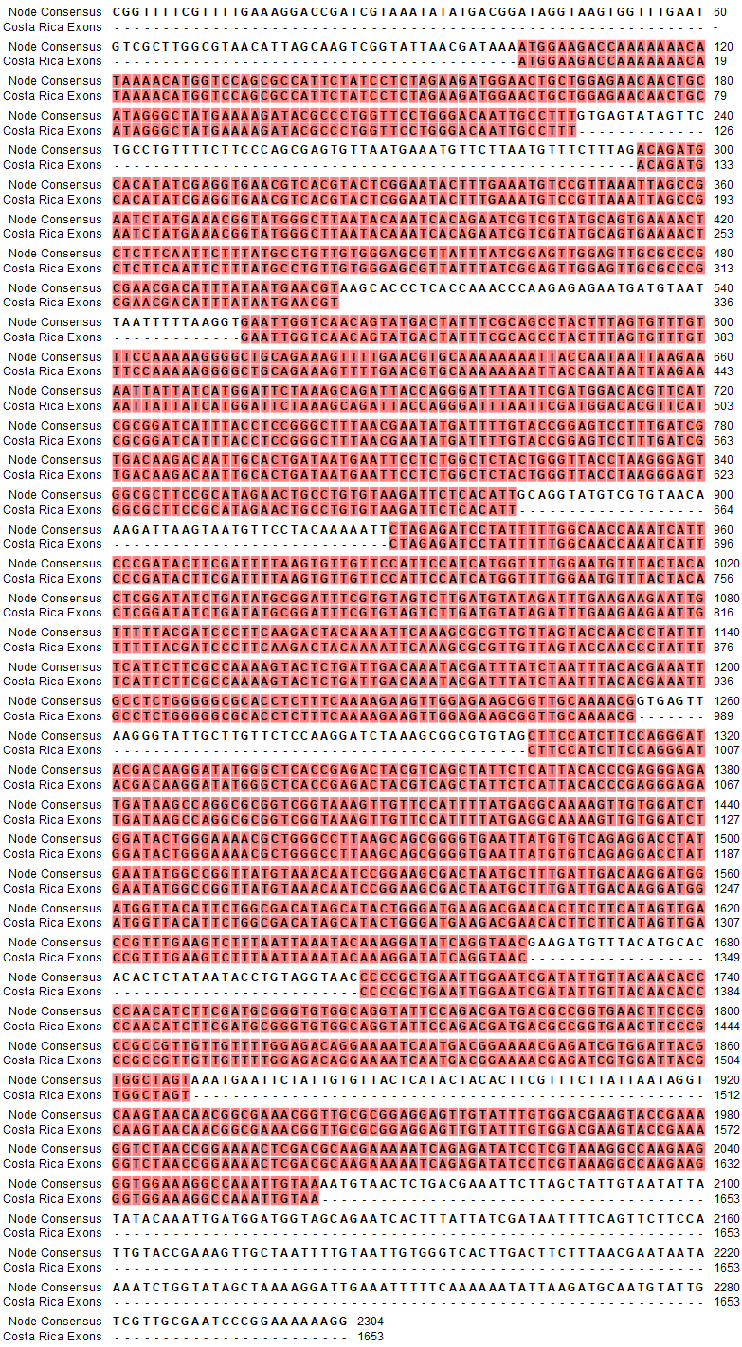

**SI Figure S9: Node consensus alignment with predicted exons for CRLuc.** The alignment of the consensus Node contigs of the Costa Rica luciferase gene with the predicted exon sequences of the Costa Rica luciferase. The seven exon regions of the two sequences are highlighted in red.

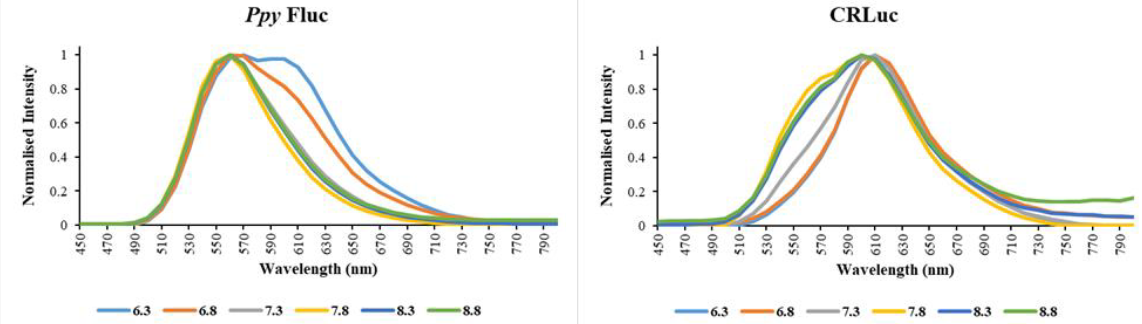

**SI Figure S10: Normalised bioluminescence spectra against pH for Ppy Fluc and CRLuc with LH2.** Luminometric measurements by substrate injection to each enzyme after 30 seconds RT incubation. Light emission from 2 second integrals every 10 nanometers from 450 to 800 nanometers. Average of triplicate values shown with data normalised with light intensity relative to lambda max.

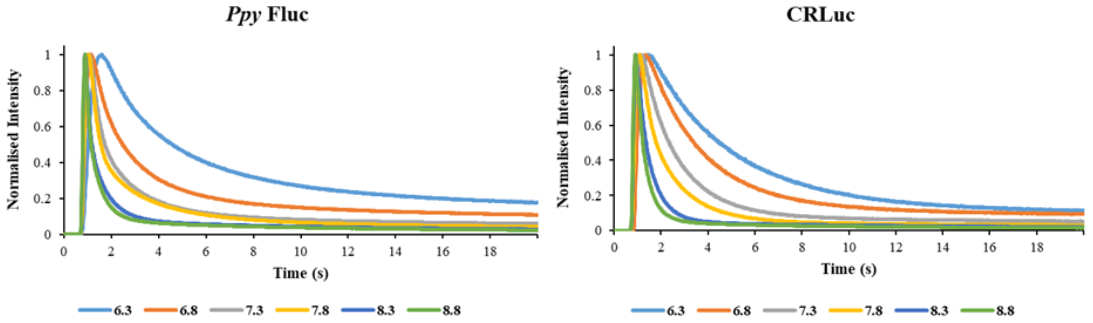

**SI Figure S11: Normalised bioluminescence against pH for Ppy Fluc and CRLuc with LH2 for 20 seconds.** Luminometric measurements by substrate injection to each enzyme. Light emission from 20 millisecond integrals for 1000 measurements. Average of triplicate values shown for each pH with data normalised with light intensity relative to emission peak.

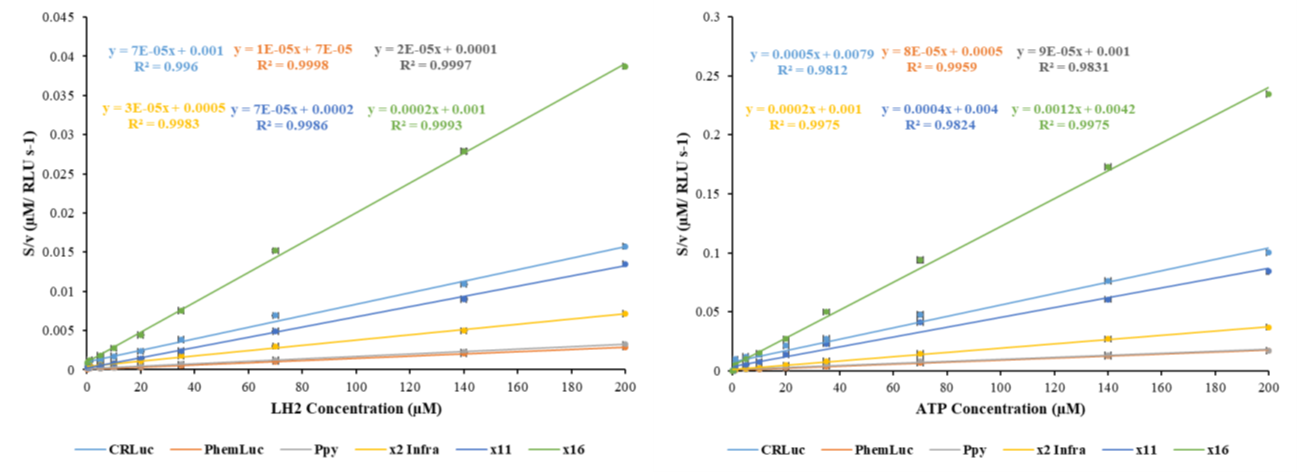

**SI Figure S12: Hanes-Woolf plots of Ppy Fluc and CRLuc with LH2 and ATP.** Plots of [S]/v against [S], where [S] is the substrate concentration and v is the estimated initial rate at that concentration indicated by Imax. Triplicate measurements of Imax over substrate concentration ranges for ATP and LH2 were used to calculate average kinetic parameters. In all assays the final concentration of enzyme was estimated to be 0.167 micromolar. For kinetic parameter calculations for LH2, assays were performed with final concentrations between 0.1 and 200 micromolar, saturated with 1 millimolar ATP. Kinetic parameters of ATP measurements, assays were performed with final concentrations between 0.1 and 1000 micromolar, saturated with 500 micromolar LH2.

sequenced

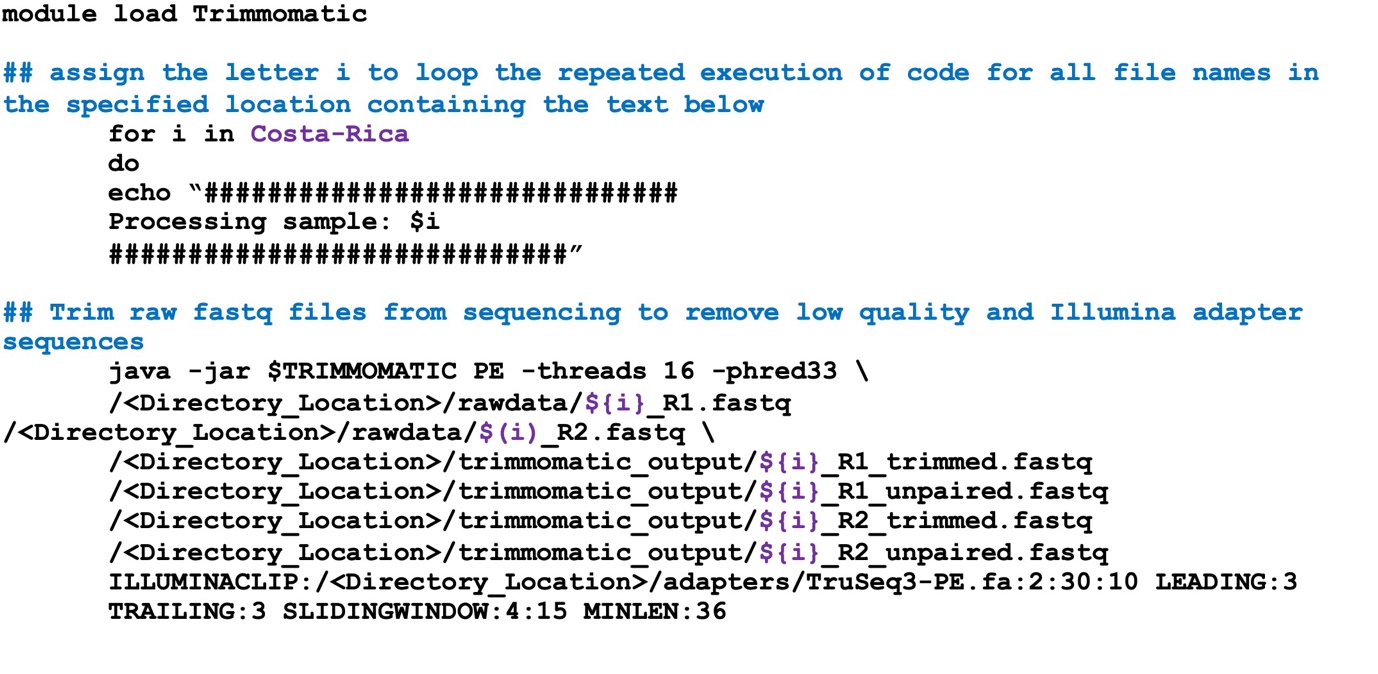

**SI Figure S13: BASH script for Trimmomatic with annotations.** Script to process sequenced libraries with Trimmomatic (Bolger *et al.* 2014). Annotations that describe the function of the following section of code are shown in blue. Sample names and their designation of ‘${i}’ are shown in purple to indicate where in the script each sample name will be substituted. The script is looped to process each sample. R1 and R2 files are forward and reverse paired reads, respectively. Singletons are reads which have no identified pairing. Trimmomatic is available at <https://github.com/usadellab/Trimmomatic>.

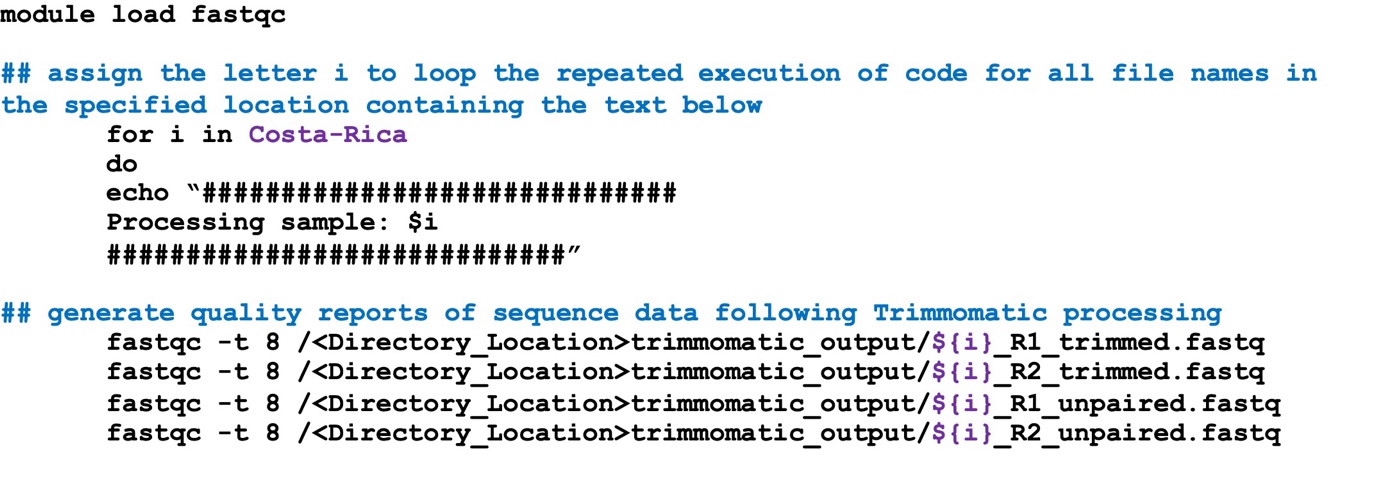

**SI Figure S14: BASH script for FastQC with annotations.** Script to generate quality reports on trimmed libraries with FastQC. Annotations that describe the function of the following section of code are shown in blue. Sample names and their designation of ‘${i}’ are shown in purple to indicate where in the script each sample name will be substituted. The script is looped to process each sample. R1 and R2 files are forward and reverse paired reads, respectively. Singletons are reads which have no identified pairing. FastQC is available at <https://www.bioinformatics.babraham.ac.uk/projects/fastqc/>.

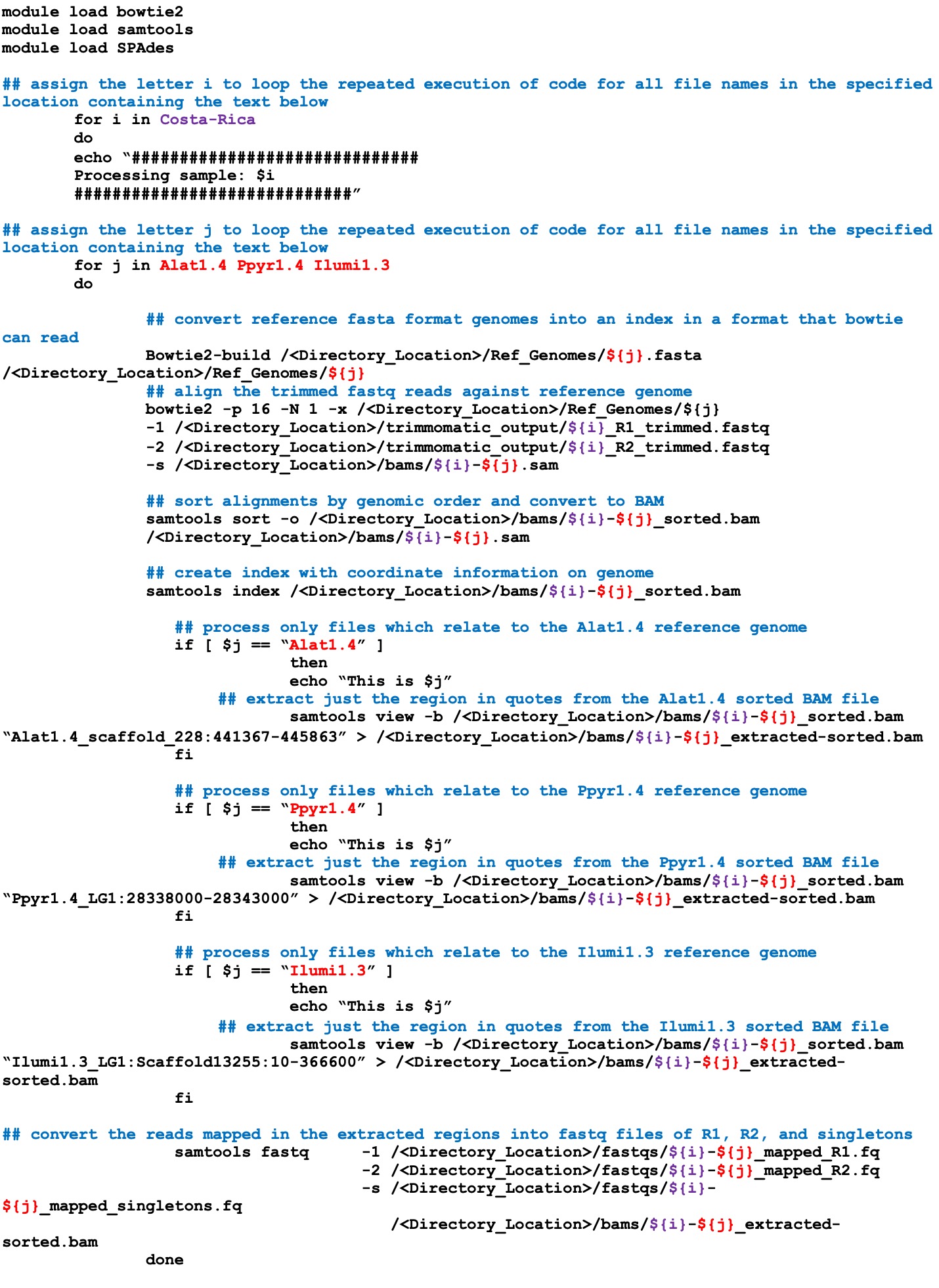

**SI Figure S15: BASH script for luciferase gene extraction with annotations.** Script to extract reads corresponding to luciferase gene sequences from trimmed libraries through the execution of Bowtie2 (Langmead and Salzberg 2012), SAMtools (Li *et al.* 2009), and SPAdes (Bankevich *et al.* 2012). Annotations that describe the function of the following section of code are shown in blue. Sample names and their designation of ‘${i}’ are shown in purple to indicate where in the script each sample name will be substituted. The script is looped to process each sample. Similarly, reference genomes and their designation of ‘${j}’ are shown in red to indicate where in the script each reference genome name will be substituted. Again, the script is looped to process each sample. R1 and R2 files are forward and reverse paired reads, respectively. Singletons are reads which have no identified pairing. Bowtie2 is available at https://sourceforge.net/projects/bowtie-bio/. SAMtools is available at http://samtools.sourceforge.net. SPAdes is available at https://github.com/ablab/spades. Firefly reference genomes Alat1.4, Ppyr1.4, and Ilumi1.3 are available at <http://fireflybase.org/>.

| **Purified and desalted Fluc** | **Ppy Fluc** | **CRLuc** |
| --- | --- | --- |
| **Size (KD)** | 60.75 | 60.71 |
| **Bradford (mg/ml)** | 0.82 | 0.057 |
| **Bradford (µM)** | 13.51 | 0.93 |
| **SDS-PAGE corrected (mg/ml)** | 0.566 | 0.057 |
| **SDS-PAGE corrected (µM)** | 9.32 | 0.93 |

**SI Table S1: Average protein concentrations of two luciferases.** The concentration of Ppy Fluc and CRLuc as determined by Bradford assay of desalted purified protein. Values for SDS-PAGE determined by ImageJ software. The protein sizes are calculated from the protein sequences. Micromolar concentrations are highlighted in grey.

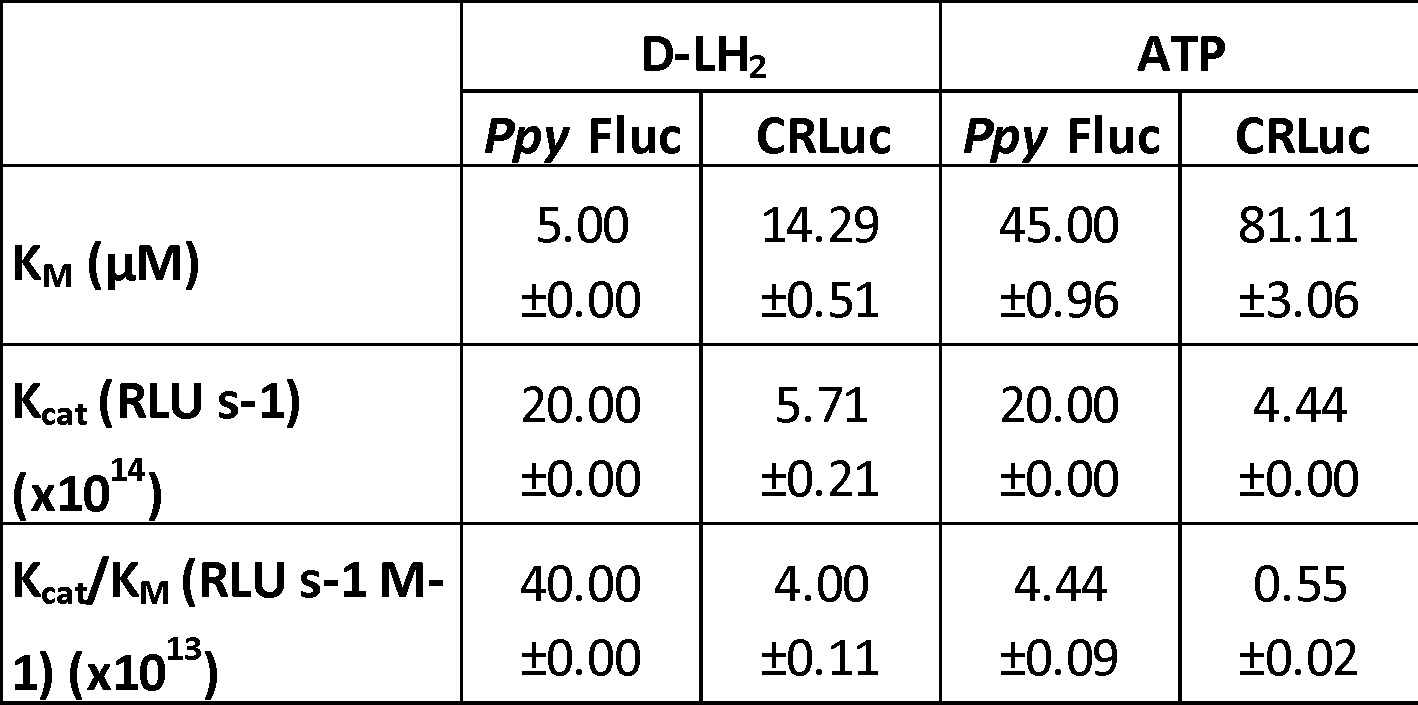

**SI Table S2: Summary of kinetic parameters for Ppy Fluc and CRLuc.** Average kinetic parameter as derived from triplicate measurements of Imax across a substrate concentration range.

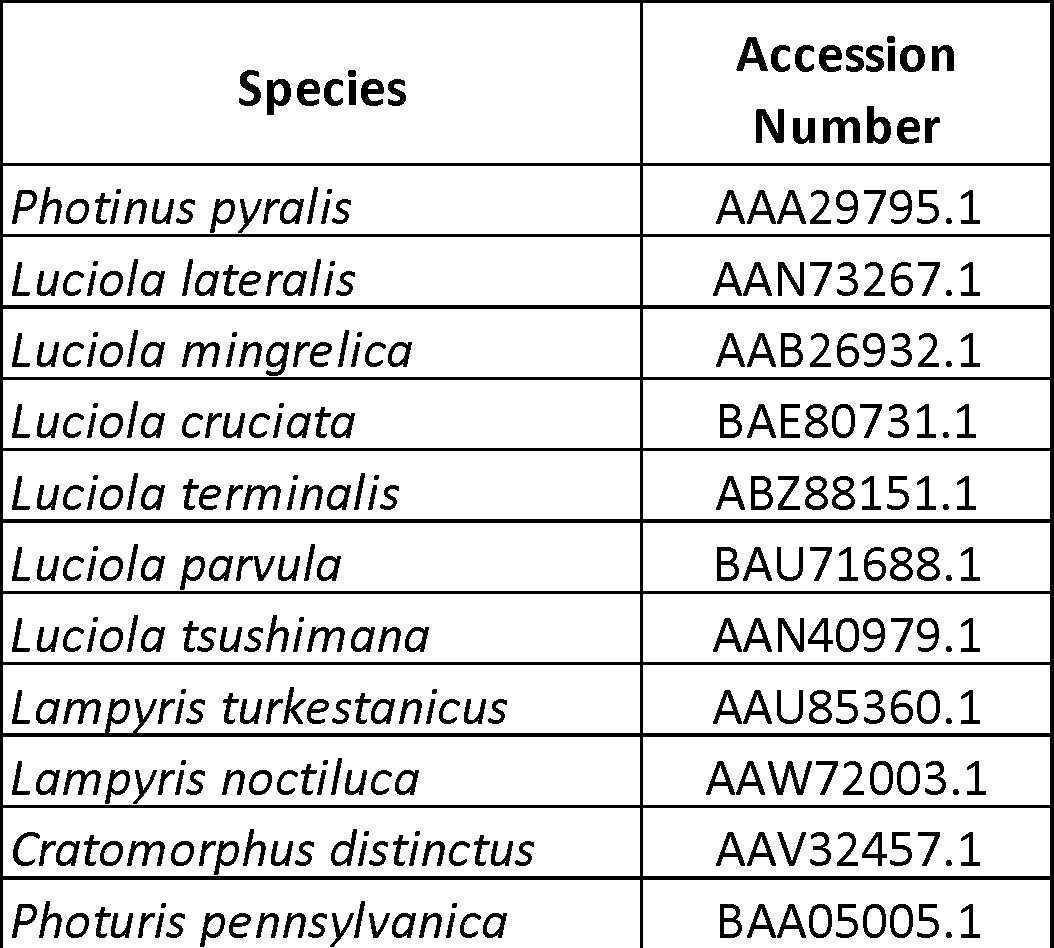

**SI Table S3: Luciferase genes used in CODEHOP design.** Eleven coleopteran luciferase genes used in the design of CODEHOP primers DKYD-F and GYG-R.

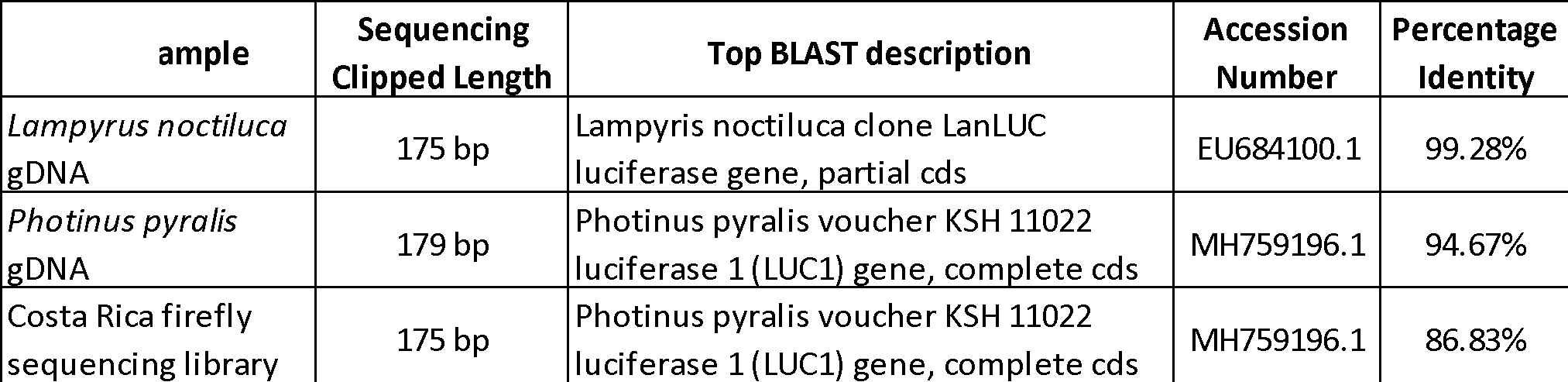

**SI Table S4: Sequencing from CODEHOP DKYD-F > GYG-R amplification.** Details of sequencing results for amplifications with CODEHOP primers DKYD-F and GYG-R. The description, accession, and percent identity of the top BLAST match are provided.

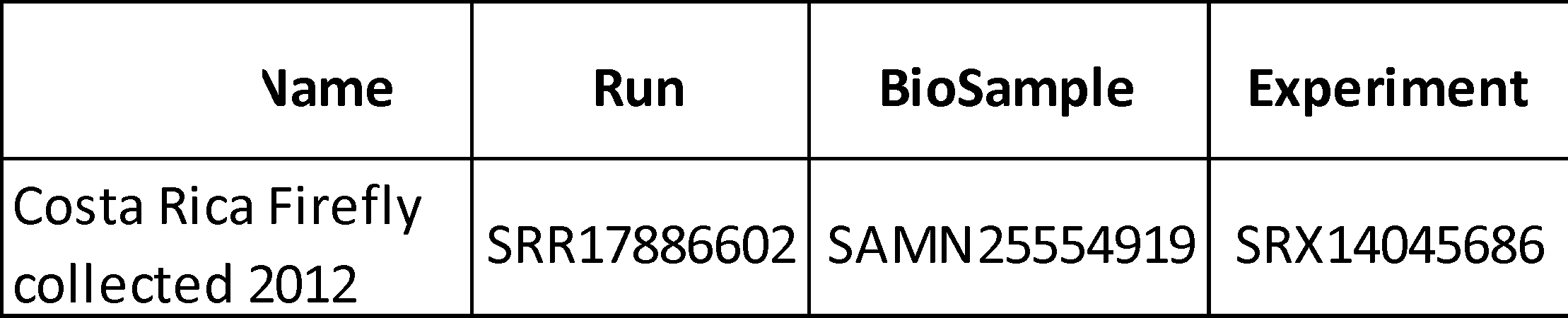

**SI Table S5: NGS data accession details.** Individual accessions of the Costa Rica firefly dataset under the BioProject accession PRJNA802557. Accessions are made available for access using the NCBI SRA Run Selector (available at https://www.ncbi.nlm.nih.gov/Traces/study/), NCBI BioSample (available at https://www.ncbi.nlm.nih.gov/biosample), and an overview of experiment details at NCBI SRA (available at https://www.ncbi.nlm.nih.gov/sra).
